# Early detection of daylengths with a feedforward circuit coregulated by circadian and diurnal cycles

**DOI:** 10.1101/2020.04.23.057711

**Authors:** Nicholas Panchy, Albrecht G. von Arnim, Tian Hong

## Abstract

Light-entrained circadian clocks confer rhythmic dynamics of cellular and molecular activities to animals and plants. These intrinsic clocks allow stable anticipations to light-dark (diel) cycles. Many genes in the model plant *Arabidopsis thaliana* are regulated by diel cycles via pathways independent of the clock, suggesting that the integration of circadian and light signals is important for the fitness of plants. Previous studies of light-clock signal integrations have focused on moderate phase adjustment of the two signals. However, dynamical features of integrations across a broad range of phases remain elusive. We recently found that phosphorylation of RIBOSOMAL PROTEIN OF THE SMALL SUBUNIT 6 (RPS6 or eS6), a ubiquitous post-translational modification across kingdoms, is influenced by the circadian clock and the light-dark (diel) cycle in an opposite manner. In order to understand this striking phenomenon and its underlying information processing capabilities, we built a mathematical model for the eS6-P control circuit. We found that the dynamics of eS6-P can be explained by a feedforward circuit with inputs from both circadian and diel cycles. Furthermore, the early-day response of this circuit with dual rhythmic inputs is sensitive to the changes in daylength, including both transient and gradual changes observed in realistic light intervals across a year, due to weather and seasons. By analyzing published gene expression data, we found that the dynamics produced by the eS6-P control circuit can be observed in the expression profiles of a large number of genes. Our work provides mechanistic insights into the complex dynamics of a ribosomal protein, and it proposes a previously underappreciated function of the circadian clock which not only prepares organisms for normal diel cycles but also helps to detect both transient and seasonal changes with a predictive power.

## Introduction

Circadian clocks provide animals, plants and certain microbes with rhythmic dynamics. These circadian pacemakers confer fitness advantages to the organisms by establishing stable anticipation of diel light-dark cycles (1, 2) as well as regulating broad morphological changes over the course of the year (3). In the plant model organism *Arabidopsis thaliana (A. thaliana)*, the circadian clock consists of at least four interacting gene modules that form negative feedback loops, which govern an intrinsic oscillator (4, 5). The expression dynamics of several hundred genes in *A. thaliana* are controlled by this circadian oscillator (6, 7). The circadian oscillator retains its essential dynamical features under constant light conditions, reflecting its intrinsic nature (8), but the phase of the oscillations can be adjusted in a process called entrainment by the lights-on signal at dawn (5, 9). In addition, however, the light-dark cycle regulates a large number of genes in a clock-independent fashion (8, 10). Therefore, molecular activities in plants are influenced by at least two periodic signals. Previous studies have found that cellular activities in *A. thaliana*, such as the abundance of the regulator of flowering time CONSTANS (CO) and the cytosolic calcium concentration, require signals from both the clock and the light-dark cycle to regulate timing (i.e. the phase of peak activity) (8, 9, 11). For example, for many mRNA transcripts, the circadian clock leads to a moderate phase shift in the onset or peak time as compared to the light cycle alone (8, 9, 11). However, these paradigms only represent *one* of the many diverse co-regulation modes of clock-light signal integration. It remains unclear whether the integration of circadian and diel light-dark cycles underpins other paradigms of signal processing.

The phosphorylation of RIBOSOMAL PROTEIN OF THE SMALL SUBUNIT 6 (RPS6 or eS6) is a post-translational modification that occurs in a wide range of organisms including animals and plants (12-14). Mice that have only a non-phosphorylatable eS6 show abnormalities at the organismal level (e.g. muscle weakness), indicating that this modification is functionally significant (15). However, while it has been suggested that the eS6 phosphorylation (eS6-P) is implicated in ribosome biogenesis in mammals (12), its biochemical consequence and specific activity, including its molecular role in *A. thaliana*, are largely unknown (13). Interestingly, eS6-P is widely used as a bioreporter for the activity of Target Of Rapamycin (TOR) kinase, a central controller of cell growth and aging (16, 17). We recently found that eS6-P in *A. thaliana* is co-regulated by both the circadian clock and the diel cycle, as eS6-P exhibits cycling behavior both in a severely clock-deficient strain and under constant light conditions (18). Unlike other extensively studied cellular processes, eS6-P is controlled by the clock and light-dark cycles in a strikingly opposite manner: in the wild-type strain under constant light the circadian clock drives eS6-P with a peak during the subjective night, whereas in a clock-deficient strain the diel light-dark cycle drives eS6 with a peak during the day (18, 19). However, the biological significance of this remarkable phenomenon at cellular and organismal levels remains elusive.

In this study, we constructed a mathematical model to examine the signaling network that regulates eS6-P in *A. thaliana*. We calibrated the model with experimental measurements of the circadian clock and eS6-P under various conditions. We found that the key dynamics of eS6-P can be explained by a feedforward loop that connects the periodic light signals to eS6-P with a direct arm and an indirect arm via the circadian clock. Although this feedforward loop is largely incoherent at the steady state of a symmetrical light-dark cycle (12-hour-light and 12-hour-dark), the amplitude of its output exhibits a high sensitivity to variations in daylength, due to the interaction between the two cyclic components in the loop. Notably, we found a characteristic early day peak of eS6-P that detects and anticipates long days. Furthermore, we combined the model with realistic photoperiod data containing both transient perturbations and long-term variations of light-dark cycles and demonstrated that the detection of daylength variations by this circuit communicates information about changes of both the season and the local environment throughout the year. By comparing our model with several representative competing models with various circuits transmitting only light signals, we found that the robust detection of day length variations requires both the circadian clock and the clock-independent light sensor. Our work demonstrates a remarkable information processing capacity of a feedforward loop that integrates circadian and light-dark cycles, and it reveals a previously underappreciated role of the circadian clock in anticipating and predicting the changes in light conditions rather than the stable photoperiod.

## Results

### A mechanistic mathematical model characterizes dynamical features of a feedforward loop controlling eS6-P

To gain a better understanding of the mechanisms underlying the dynamics of eS6-P, we built a mathematical model that includes a light-entrained circadian clock and a light-dependent, clock-independent signaling pathway that regulates the phosphorylation of eS6 (Fig 1A). The latter light-pathway consists of TOR, a kinase that transmits light signals, and ribosomal S6 kinase (S6K), a substrate of TOR (20-23). S6K has been shown to phosphorylate eS6, and the TOR-S6K axis is known as a primary pathway for eS6-P in multiple organisms including plants (17). The clock component of the model is described by four interacting components (LHY/CCA1, the evening complex (EC), PRR9/PRR7, and PRR5/PRR1, denoted as C1, C2, C3 and C4 respectively in this study). It contains the core repressilator of the plant circadian circuit and has previously been used to model the circadian clock (Fig 1A) (4, 24-26). We used this model to capture the essential dynamical features of the clock rather than the molecular details, so we focused on this four-component core repressilator and neglected the additional feedbacks in the clock (4). Because we previously observed that eS6-P oscillates under constant light (18), we also considered clock-driven regulation of eS6 phosphorylation and dephosphorylation in our model (Fig. 1A). Before parameter estimation, each clock component in the model was assumed to regulate (activate or inhibit) the phosphorylation or dephosphorylation of eS6 with unknown parameters. Because we assumed that the clock does not receive signals from the clock-independent pathway or eS6, we first obtained a parameter set for the clock component by fitting the clock model using a qualitative objective function to capture the basic behaviors including cyclic variation in response to light-dark cycles (entrainment) and cycling under constant light conditions (see Methods, Fig. S1). We then compared the simulated clock gene dynamics to previously published expression data (27) to ensure that the model properly represents the phase and shape of the time course measurements (Fig. S2).

**Fig. 1.**
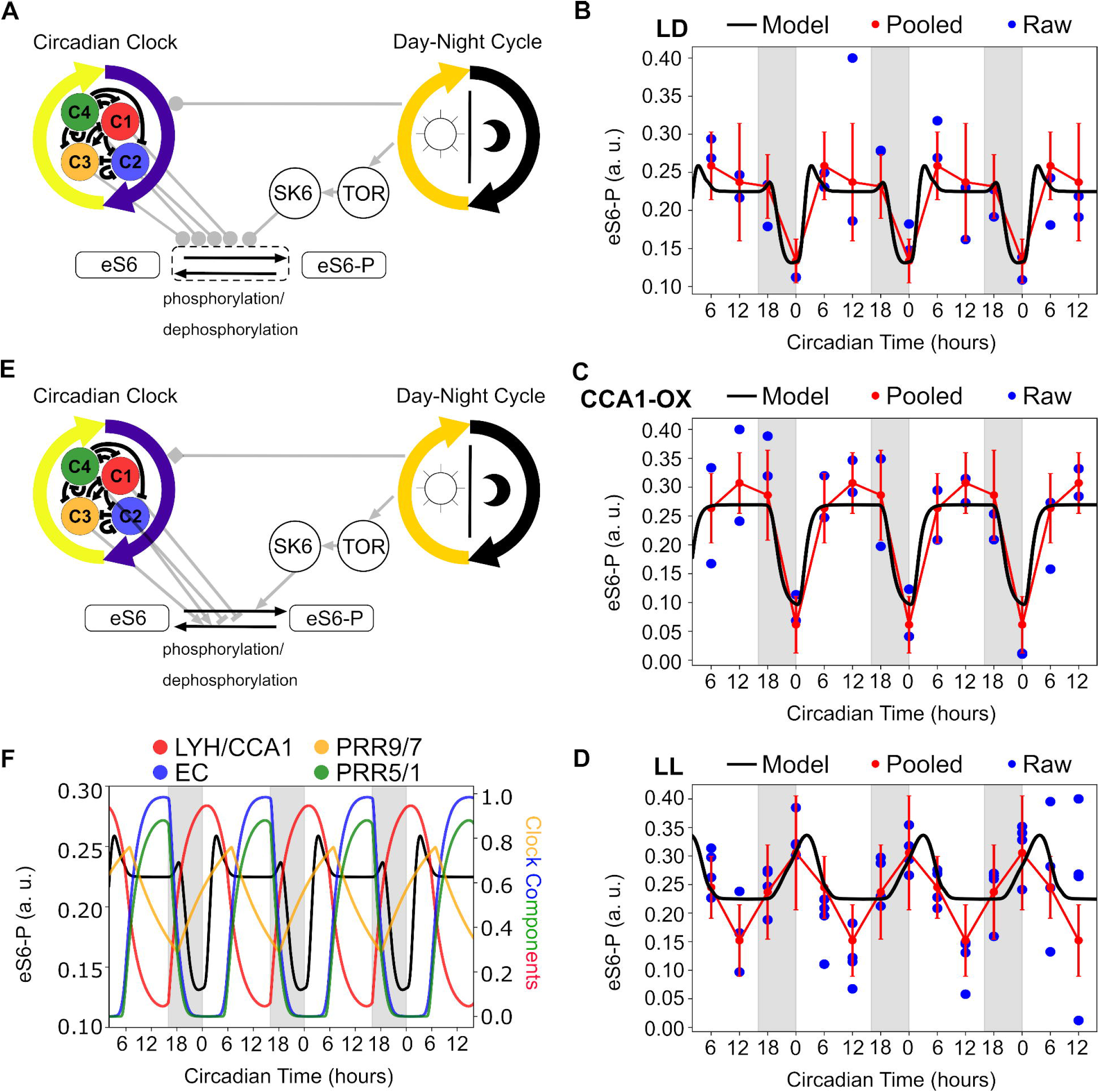
Network structure of the eS6-P regulatory circuit and comparisons between simulations and experiments. **(A)** An influence diagram of a model of the light and clock driven feedforward regulatory system that regulates eS6-P (Clock+Light Model). This model consists of two cycling systems: the light-dark cycle (an oscillatory input) and the circadian clock (an autonomous oscillator). Each cycle regulates cellular processes independently. The regulation by light-dark cycle is mediated by TOR-S6K pathway, and the clock driven regulation is mediated by transcription factors belonging to the LHY/CCA1 (C1), Evening Complex (EC,C2), PRR9/7 (C3), PRR5/1 (C4) modules which contain a repressilator circuit and additional interactions. The light-dark cycle also regulates the circadian clock via the LHY/CCA1 and PRR9/7 modules (entrainment), thus creating a feedforward circuit. Arrows with circle head show influence of unknown directions (positive or negative) before parameter estimation. **(B)** Model predictions (black) and experimental observations (red) of eS6-P under wild-type in long-days (16:8, LD). Error bars indicate standard deviation of pooled measurements at various circadian times. Circadian time is relative to dawn, and regions shaded with grey indicate period of darkness. Blue dots show the raw data points collected prior to pooling over a period of 84 hours (18). **(C)** Model predictions (black) and experimental observations (red) of eS6-P under long-days (16:8, LD) but with a deficient clock (CCA1-overexpression). Error bars indicate standard deviation of pooled measurements at various circadian times. Time is measured and the graph is shaded as in **(B). (D)** Model predictions (black) and experimental observations (red) of eS6-P under constant light. Error bars indicates standard deviation of pooled measurements at various circadian times. Circadian time is measured relative to subjective dawn (Zeitgeber time, ZT) and regions shaded grey indicate periods of subjective night. eS6-P is shown in arbitrary units (a. u.) (see Methods). **(E)** An influence diagram of our model of eS6-P updated with the direction (triangle arrowhead: activation, flat arrowhead: repression). Note that the ambiguous (diamond) regulation of the clock by light reflects that light regulates LHY/CCA1 and PRR9/PRR7 in opposite directions (we assume that light represses CCA1 stability [49], which is important to restrict the LHY/CCA1 peak to dawn). **(F)** An overlay of clock element activities on top of the eS6-P trajectory in wild type under a long-day cycle. The eS6-P cycle is indicated by the solid black curve and the clock components are represented by colored curves (red = C1 = LHY/CCA1, blue = C2 = EC, orange = C3 = PRR9/PRR7, green = C4 = PRR5/PRR1). Circadian time is measured relative to dawn.

Next, we fit the Clock+Light model that contains both clock-dependent and clock-independent pathways regulating eS6-P to our recent measurement of eS6-P dynamics under three experimental conditions: a wild-type (WT) strain under long day (LD, 16-hour-light and 8-hour-dark) condition, the WT strain under constant light (LL) condition, and a clock-deficient strain under LD condition (CCA1-overexpression) (18). Using an evolutionary algorithm with a likelihood-based objective function, we found optimized parameter sets that produced trajectories that reasonably matched the experimental data in all three conditions (Fig. 1B-D). Importantly, the model captured the remarkable dynamical features of eS6-P: light alone drives the upregulation of eS6-P during the day (Fig. 1C) whereas the clock alone drives the upregulation of eS6-P during the subjective night (Fig. 1D). Our model also recapitulated the day-peak of the eS6-P when both clock and light-dark cycles are present, suggesting the dominant role of the light-dark cycle. Furthermore, the model predicted a peak of eS6-P shortly after dawn that was not measured experimentally. We found that this observation was not due to the choice of a particular parameter set: the majority of the top 70 (or 5%) models (based on likelihood) from multiple optimization runs generated the same behavior (Fig. S3A). Conversely, the next 70 models lacked the dawn peak, but those models also lacked oscillations under constant light, suggesting a loss of clock regulation (Fig. S3B). Our subsequent analyses are based on the top performing model, which produced a moderate peak of eS6-P after dawn.

We next examined how the clock and the light-dark cycles influenced eS6-P mechanistically. While the clock-independent pathway had an obvious mechanism of action, in which the light signal activates a cascade of two kinases that give rise to eS6 phosphorylation, the clock pathway involved a nontrivial combination of molecular influences. We found that eS6-P is regulated by multiple clock-genes that peak at different time points during the day (Fig 1E). Specifically, LHY/CCA1 and PRR9/PRR7 both regulated eS6 during the early day with different phases of activity (LHY/CCA1 at dawn and PRR9/PRR7 a few hours after dawn) and opposite effects on eS6 dephosphorylation (LHY/CCA1 inhibits and PRR9/PRR7 promotes dephosphorylation). In contrast, while EC and PRR5/PRR1 also have opposing influences, their activities largely overlapped and canceled each other out. These clock influences collectively resulted in the stably high amount of eS6-P between the early day and dusk, while the early day peak resulted from a coincidence of light, the peak LHY/CCA1 activity, and the absence of peak PRR9/PRR7 activity (Fig 1F). As such, our model showed that at the steady state under LD, the circadian clock and the light-dark cycle influenced eS6-P in a generally opposing manner: during the day, eS6-P is promoted by light but inhibited by PRRs, while during the night, the anticipatory rise of LHY/CCA1 promoted eS6-P prior to dawn, leading both to rapid early rise and the early day peak under normal LD conditions, and the shift in the rise into the subjective night under constant light. Considering the clock’s dynamics are light-entrained, the resulting model behaved as a feedforward loop that is largely incoherent at steady state.

In conclusion, our calibrated model captured the observed dynamics of eS6-P under multiple conditions, and allowed us to make predictions about the mechanisms underlying these intriguing dynamics. Note that although our discussion primarily focuses on one optimized model, we have reproduced our key conclusions with distributions of parameters rather than a single parameter set (Fig. S3), and with a much more detailed clock model which includes both mRNA and protein concentrations (Fig. S4) (4).

### Integration of circadian clock and light-sensing pathways detects and anticipates long daylength upon transient perturbations of light-dark cycles

We next focused on the dynamics of eS6-P after dawn, when the coincidence of clock and light signals gave rise to a sharp rise and a peak in the model (Fig. 1B-D). We hypothesized that at this key time interval a change in the phase of the lights-on signal significantly affect the dynamics of the model output because the distinct influences of light and clock can synergize with each other depending on their relative phase. We therefore focused on variations in the timing of the transition from night to day (t_ND_), as they might be used as proxies for transient changes in the local environment (such as weather)or long-term seasonal changes. We ran simulations with the optimized Clock+Light model (Fig. 1E) under the 12-hour-light and 12-hour-dark (12L:12D) condition. After the system reached steady state (Day 0), we varied t_ND_ (+/- 4 hours) at the start of a single day (Day 1), and tracked the trajectories of eS6-P from the perturbed day and onwards (Fig. 2A). We found that the transient variation of t_ND_ resulted in significant changes in the abundance of eS6-P (response) in the early day, but the responses converged as the day continued. In particular, an early night-to-day transition time (t_ND_<0) gave rise to a higher response including an early day peak, as previously observed with the base model under LD condition (Fig. 1B, Fig. 2B purple), whereas a late transition time (t_ND_>0) had the opposite effect and resulted in a trough (Fig. 2B yellow). In addition, early day maximum eS6-P (defined as the maximum eS6-P over the 4 hours after dawn) varied by as much as 13.8% relative to the case of Δt_ND_ = 0. We found that the early day peaks (transient responses that are greater than the steady-state response during the day) appeared only when the daylength exceeded 12 hours (Fig. 2B cyan). These results show that the eS6-P control circuit detected phase variations of the lights-on signal and thus effectively sensed daylength variation at the beginning of the day.

**Fig. 2.**
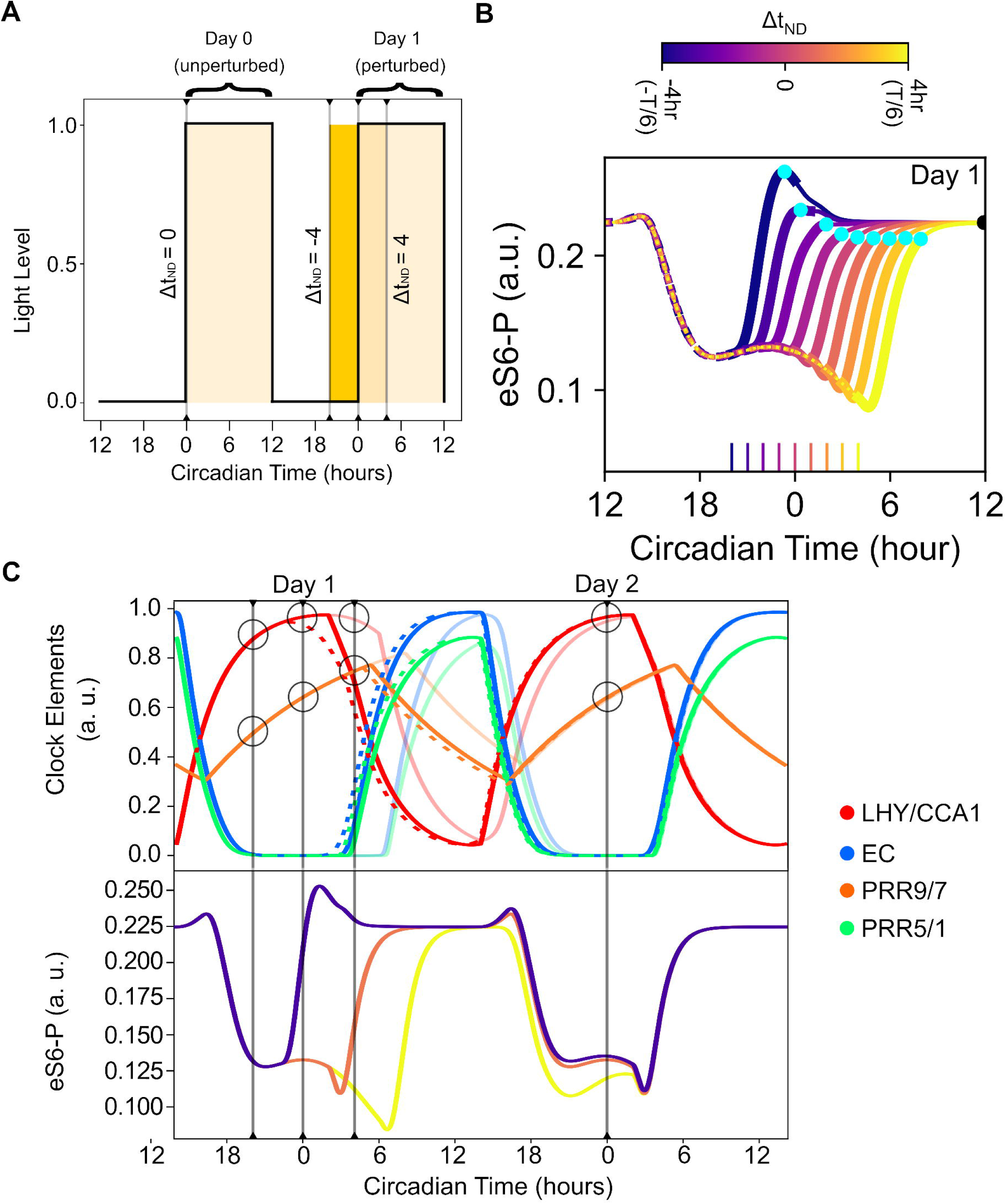
Response of eS6-P to variations of the night-to-day transition time. **(A)** Diagram of perturbations of the night-to-day transition and the effect of daylength relative to a 12-hour-light-12-hour-dark (12L:12D) day. The unperturbed (Δt_ND_ = 0) and perturbed (Δt_ND_ +/- 4 hours) days are labeled. The day period is shown by yellow shaded regions and the maximum extent of the change in daylength is shown by differential shading. The system reached steady state before Day 0. **(B)** Trajectories of eS6-P in response to variations of the night-to-day transition time from Δt_ND_ = -4 (purple) to Δt_ND_ = 4 (yellow) in 0.5-hour increments. All trajectories start at the dusk of the previous day (Day 0, unperturbed), and end at the dusk of the current day (Day 1, perturbed). Dotted lines show trajectories in the absence of light. Thick solid curves show trajectories in the early day (first four hours after dawn), and thin solid curves show trajectories in the remaining hours of the day. Short line segments at the bottom show the time of dawn for each trajectory. **(C)** Side by side plots of clock perturbations in response to varying night to day transition and the resulting effects on eS6-P. For simplicity, we only show the Δt_ND_ = -4 (purple), Δt_ND_ = 0 (red), Δt_ND_ = 4 (yellow) in the bottom figure. In the top panel, color indicates clock factor (red = C1 = LHY/CCA1, blue = C2 = EC, orange = C3 = PRR9/PRR7, green = C4 = PRR5/PRR1), while shade of color represents the timing of dawn (dashed, Δt_ND_ = -4; regular solid, Δt_ND_ = 0; semitransparent solid, Δt_ND_ = 4). Dashed lines show the corresponding times while circles on the clock factors’ curves highlight how the ratio between LHY/CCA1 and PRR9/PRR7 corresponds to peak height of the response. In all panels, Circadian time is shown in hours relative to normal time of dawn (i.e. +/- 0 hours).

We found that the sensitivity of eS6-P to changes in the light-dark cycle was associated with the altered relative phase of clock-gene activities with respect to the dawn (Fig. 2C, gray lines) upon perturbations of the light-dark cycle. For example, the earlier dawn allowed synergy between light signal, peak of CCA1/LHY and relatively low activities of PRR modules to give rise to a strong eS6-P response (Fig. 2C). This change of relative phase occurred despite the influence of the transient phase shift on the clock gene dynamics as a form of entrainment (Fig. 2C upper panel). Variation in photoperiod length has been previously shown to affect the absolute phase of circadian genes, altering both their timing relative to dawn and the intervals between peak expression of different circadian genes, with the gap between CCA1/LHY and PRR9 growing as the photoperiod becomes longer (26, 28). These results suggest that the circadian clock plays an essential role in the regulation of eS6-P, particularly with regard to its early day dynamics.

We therefore asked whether the circadian clock is required for the eS6-P circuit (Fig. 1E, Clock+Light model) to detect long daylength with an early day peak. To this end, we constructed three alternative models that describe various modes in which eS6-P may respond to the light signal in the absence of the clock (Fig. 3A, see Table S2 for parameters). These models are: 1) a linear circuit that transmits light signals to eS6-P (Fig. 3A, top panel), 2) an incoherent feedforward loop (IFFL) that produces an early day peak similar to what the Clock+Light model does under LD condition (Fig. 3A and B, middle panels), and 3) a coherent feedforward loop (CFFL) that allows slow decline of eS6-P upon the withdrawal of light signals (Fig. 3A, lower panel). Note that none of these three models were able to fully fit the observed data (e.g. Fig. 1D) due to the lack of clock regulation. Rather than evaluating these models in terms of their fit to data, we focused on their performance in terms of detecting daylength variations upon dawn.

**Fig. 3.**
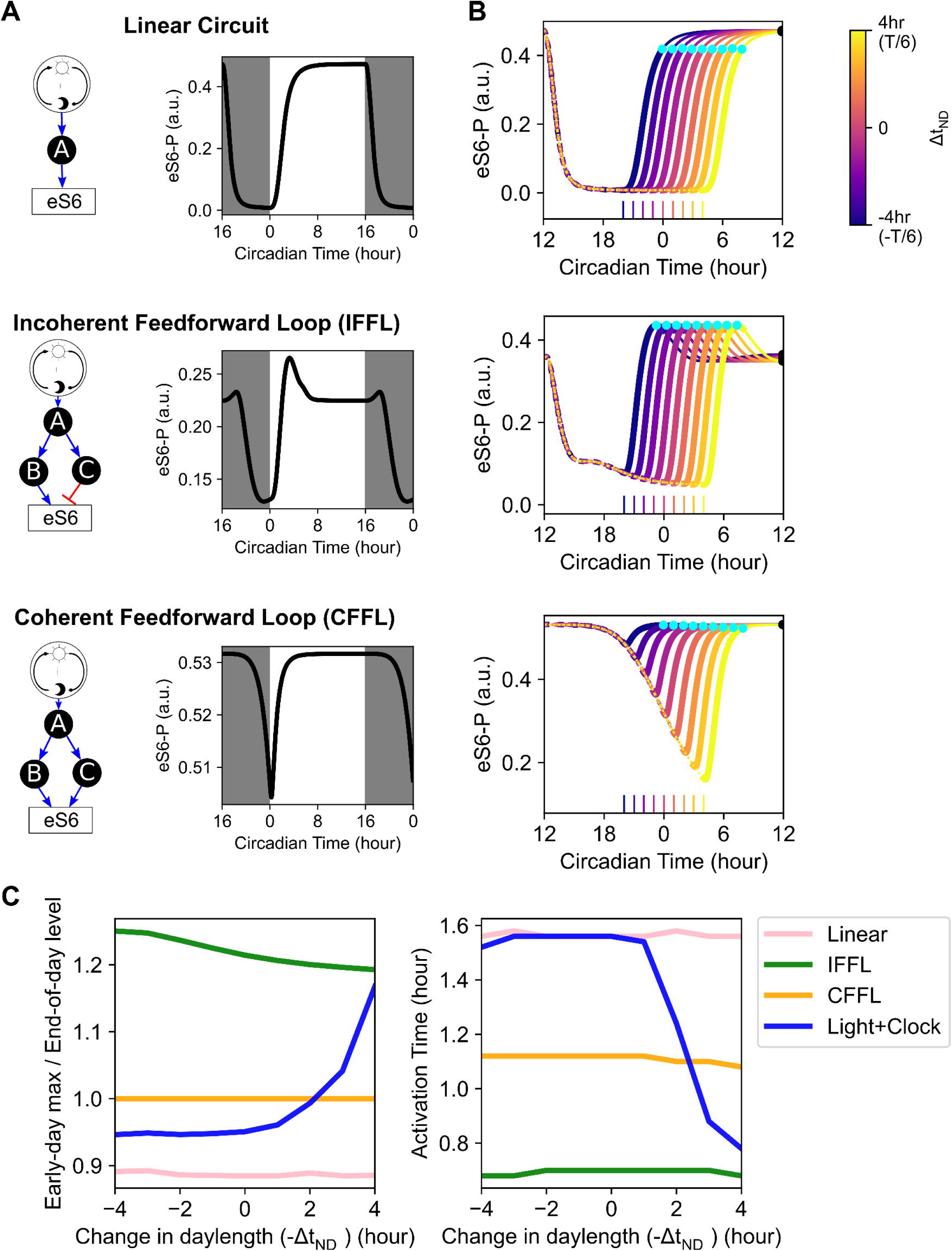
Response of alternative models of eS6-P circuit to variations of the night to day transition time. **(A)** Influence diagrams of the three alternative models of eS6-P circuit. In each diagram, the regulatory factors are indicated by the lettered black circles, and regulatory interactions are denoted by colored lines (blue = activation, red = repression). Right panels show simulation trajectories with these models under the LD condition. Gray regions show the period of night. **(B)** Trajectories of eS6-P in response to variations of the night to day transition time from Δt_ND_ = -4 (purple) to Δt_ND_ = 4 (yellow) in 0.5-hour increments. All trajectories start at the dusk of the previous day (Day 0, unperturbed), and end at the dusk of the current day (Day 1, perturbed). Dotted lines show trajectories in the absence of light. Thick solid curves show trajectories in the early day (first four hours after dawn), and thin solid curves show trajectories in the remaining hours of the day. Short line segments at the bottom show the time of dawn for each trajectory. **(C)** Left: peak metric (ratio of the early day maximum of eS6-P to the eS6-P levels at the dusk) with respect to the perturbations of the daylength. Right: activation time of the eS6-P after dawn with respect to the perturbations of the daylength. Activation time is defined as the time for eS6-P to reach *x*_0_ + 0.9(*x*_*s*_ − *x*_0_), where *x*_0_ is the eS6-P level at the dawn and *x*_*s*_ is the eS6-P level at the dusk.

Each of the three alternative models generated a constant level of eS6-P upon the perturbations of daylength (Fig. 3B cyan). Particularly, the early day peak produced by the IFFL model did not distinguish short and long daylengths (Fig. 3B middle). This insensitivity to daylength variations with the alternative models was reflected in the stable maximum early responses of eS6-P regardless of their ratios to steady state (or end-of-day) eS6-P levels (Fig. 3C left). In addition, we found that the daylength-sensitive early response with the Clock+Light model was anticorrelated with the time for eS6-P to reach its steady state level, e.g. an early dawn accelerated the response of eS6-P (Fig. 3C right). These results suggest that detection of long daylength with early day peaks is a feature that requires the integration of the circadian clock and the clock-independent light-sensing pathway.

Because realistic perturbations of daylength may occur through changes of light conditions at both dusk and dawn and on multiple days, we next considered the effects of multiple variations in our simulations. We first allowed -/+ 2 hours of variation in t_ND_ followed by +/-2 hours of variation in the day-to-night transition (t_DN_) during dusk the previous day (Fig. 4A). Perturbations in each t_DN_-t_ND_ pair acted in the same direction to either lengthen or shorten the intervening night (Fig. 4A and B, vertical lines). With the Clock+Light model, we found that consecutive changes in dusk and dawn timing gave rise to significant variability in eS6-P response, similar to what was obtained with -/+ 4 hours of t_ND_ alone (Fig. 2B, Fig. 4B and C). Interestingly, +/- 2 hours perturbations of t_ND_ or t_DN_ alone did not generate an early day peak (Fig. 4B and C). This suggests that changes in the timing of dusk can be integrated into the eS6-P response by the Clock+Light circuit after dawn and serve as a predicting factor for the current daylength. The incorporation of information in the previous day shows that the Clock+Light circuit can anticipate the daylength changes with a memory capacity.

**Fig. 4.**
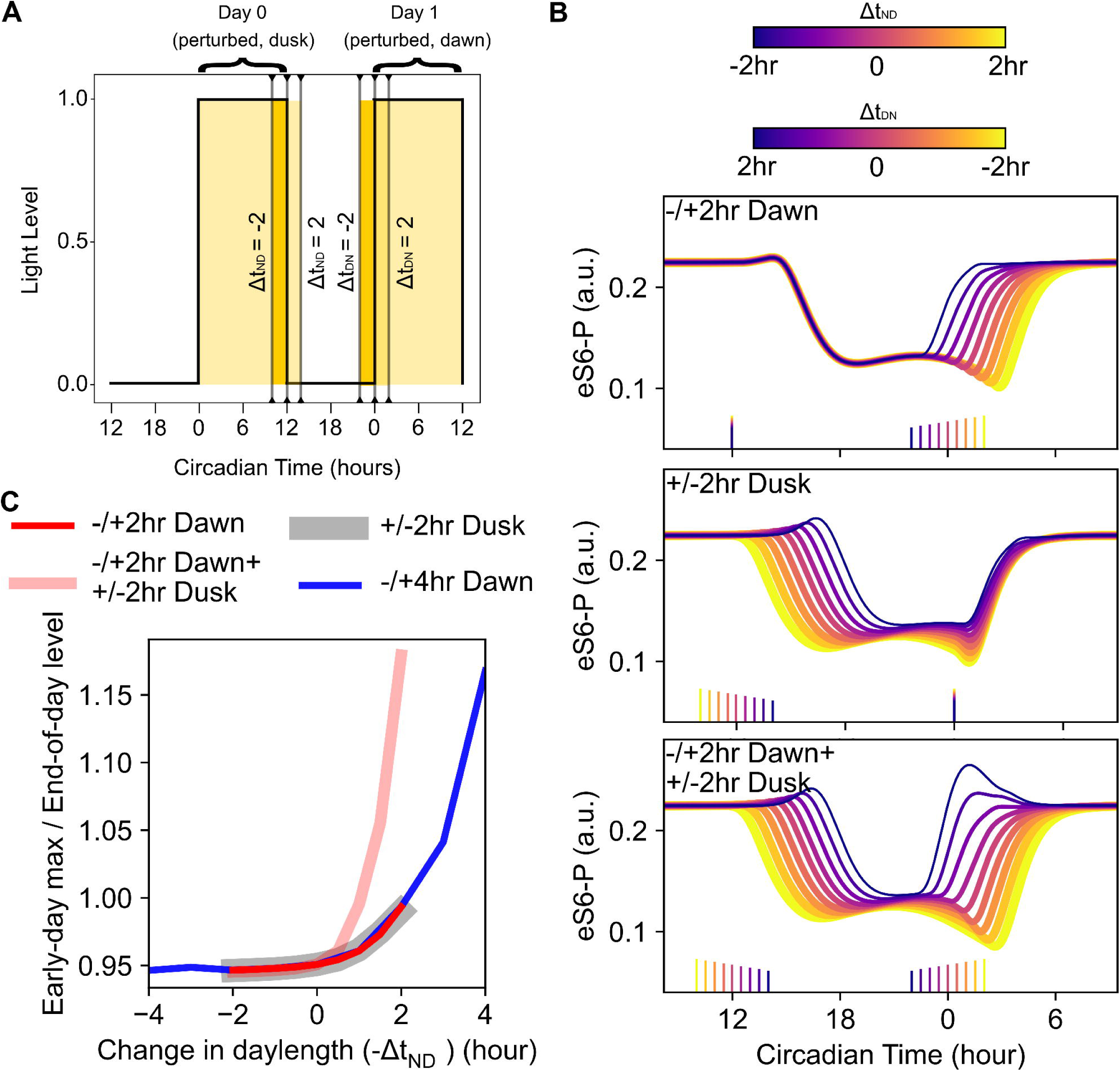
Response of eS6-P to variations in night to day transition time following changes of day to night transition time the previous dusk. **(A)** Diagram of perturbations of the day to night transition and the night to day transition on two consecutive days. The first (perturbed at dusk, Δt_ND_ = -/+ 2 hours) and second days (perturbed at dawn, Δt_ND_ = +/- 2 hours) are labeled. The day period with light is shown by yellow shaded regions and the maximum extent of the change in daylength is shown by differential shading. **(B)** Trajectories of eS6-P in response to variations of the day to night transition time and/or the night to day transition time from Δt_ND_ = -2, Δt_DN_ = 2 (purple) to Δt_ND_ = 2, Δt_DN_ = -2 (yellow) in 0.25-hour increments. Short line segments at the bottom show the time of dawn or dusk for each trajectory. Upper: dawn perturbation only. Middle: dusk perturbation only. Bottom: both perturbations. **(C)** Peak metric (ratio of the early day (Day 1) maximum of eS6-P to the eS6-P levels at the dusk) with respect to the perturbations of the daylength under various perturbation scenarios.

We next examined the effect of changes to t_ND_ on two consecutive days. We observed a slightly increased dynamic range of early eS6-P peak on the second early day compared to the first day with the Clock+Light model (Fig. S5). These results indicate that the Clock+Light circuit was able to detect changes in daylength beyond the altered timing of the current dawn.

### Integration of circadian clock and light-sensing pathway detects gradual variations of light-dark cycles

We next asked whether the eS6-P circuit responds to progressive, long-term variations of the light-dark cycle, reflective of seasonal changes in the cycle throughout a year. We used a Solar Calculator provided by the National Oceanic and Atmospheric Administration (NOAA) to generate a data set of ‘ideal’ (i.e. unaffected by transient local changes) daylengths over the course of a year. We simulated the Clock+Light model based on these light-dark cycles for one year at two locations representing the extreme latitudes of the *A. thaliana* distribution, Oslo, Norway, and Praia, Cape Verde (29). As expected from the analysis of transient changes in dawn timing, our model showed a sensitivity of eS6-P to changes in daylength over the year (Fig. 5). The range of eS6-P was correlated with the degree of variation in daylength, which was broader in Oslo (6.2 to 19.0 hours, Fig. 5A) compared to Praia (11.3 to 13.0 hours, Fig. 5B). We observed variations in both the early-day maximum and early-day minimum levels of eS6-P in Oslo (Fig. 5A, orange and pink), with the early-day maxima increasing dramatically as daylength approached its yearly maximum in the summer. Starting from Day 89 of the year, the early-day maxima increased until it reached 148% of the daily steady state at Day 170. As such, we conclude that changes in the daylength over a year are able to the variations in the early-day peak of eS6-P, with the early day maxima exceeding the daily steady state level around 13.5 hours of light. In contrast, much smaller variations of the early-day eS6-P, which never exceeded the daily steady state levels, were observed with the yearly light-dark cycle data in Praia (Fig. 5B), which has a maximum daylength of 13.0 hours. These results suggest that the eS6-P circuit can detect seasonal changes of the light-dark cycles by varying the magnitude of the responses after the dawn and thus is sensitive to changes in daylength even when they occur gradually.

**Fig. 5.**
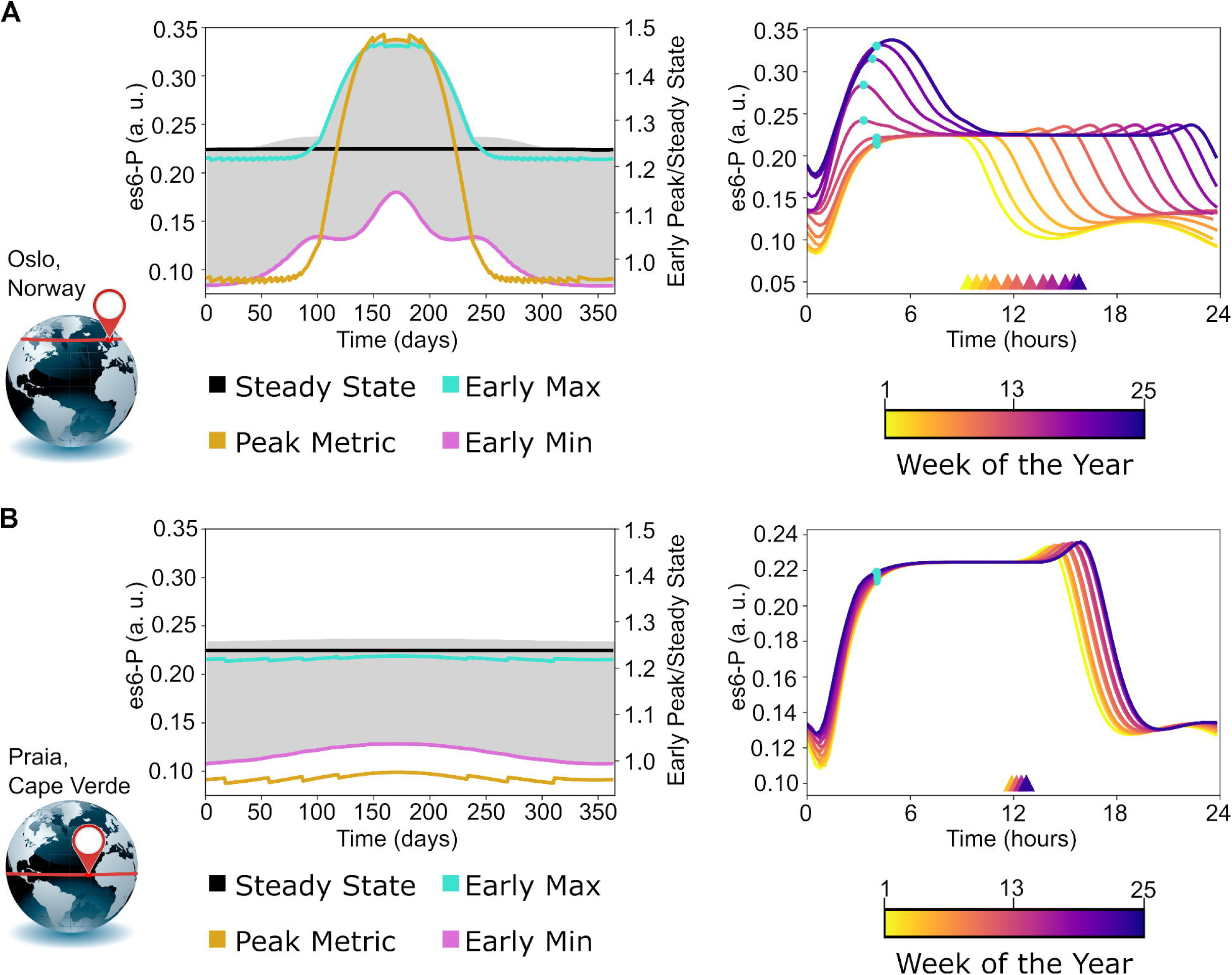
Simulated amounts of eS6-P in response to seasonal changes in daylength over a year. **(A)** Simulation of eS6-P over a full year using daylength data for Oslo, Norway (the NOAA Solar Calculator). The left figure shows the eS6-P time course over the full year (gray), overlaid with the early-day maximum (green), early-day minimum (pink), and daily steady state (end-of-day, or dusk) level (black). Peak metric (orange) is defined as the ratio of early-day maximum to the daily steady state level. The right figure shows the eS6-P profiles of individual days selected every two weeks from Day 7 to Day 175 of the year. Time 0 is the actual dawn of each day. The color of each curve corresponds to the week number and is correlated with the daylength from short (yellow) to long (purple). Triangles indicate the time at which the end-of-day eS6-P level was measured, and blue dots indicate the position of the early-day peak. **(B)** Simulation of eS6-P across a full year using daylength data for Praia, Cape Verde (the NOAA Solar Calculator). Upper and lower figures are as described in **(A)**. The globe on the left of each panel shows the latitude of the position that the daylength was simulated for.

We next asked whether the eS6-P circuit can detect changes in light-dark cycles due to transient changes (such as weather) as well as seasonal changes. We obtained a year-long environmental radiometry data from Harvard Forest (30). We normalized the measurement of downward photosynthetic radiation based on the ideal daylength calculations for Boston, Massachusetts from NOAA and obtained a time-series data of realistic light-dark cycles over a year (see Methods). Briefly, a threshold value was used to define day/night with the Harvard Forest data, and the value was chosen to minimize the mean difference between the inferred hours of daylight and NOAA estimations of daylength. We compared the simulation results for idealized and observed daylength data (Fig. 6) With the idealized daylengths in Boston (9.4 to 15.4 hours), we observed moderate early day peaks in the middle of the year (Fig. 6A). However, transient changes in the observed light data resulted in changes in daylength of up to 1.5 hours, which in turn gave rise to significant variation of early-day peak of eS6-P in response to the changes of sunlight due to daily weather in addition to seasonal changes of sunlight (Fig. 6B). Although transient changes mainly reduced the daylengths from the idealized day lengths throughout the year, there is a significant difference in terms of peak metric between longer (>13 hours) and shorter (<13 hours) days based on the realistic sunlight data (1.01 vs. 0.96. Welch’s t-test, *p* =2.7×10^−34^). Furthermore, the absolute difference in early-day peak between idealized and realistic days was greater during longer days (> 13 hours) than during shorter days (< 13 hours) by 10-fold (Welch’s t-test, *p* =7.5×10^−32^). As such, these weather induced changes primarily exist in long days, during which the early response of eS6-P is most sensitive to daylength changes. To further illustrate this point, we compared in simulated early day peaks between idealized and realistic days: in both simulations the peak to steady state ratio remained low during short days and increased around a daylength of 13 hours, but the ratios of the two simulations diverged as days grew longer (Fig. 6C). Together, these results show that the eS6-P circuit can detect variations in light-dark cycles when both seasonal and local, transient (weather) changes are considered.

**Fig. 6.**
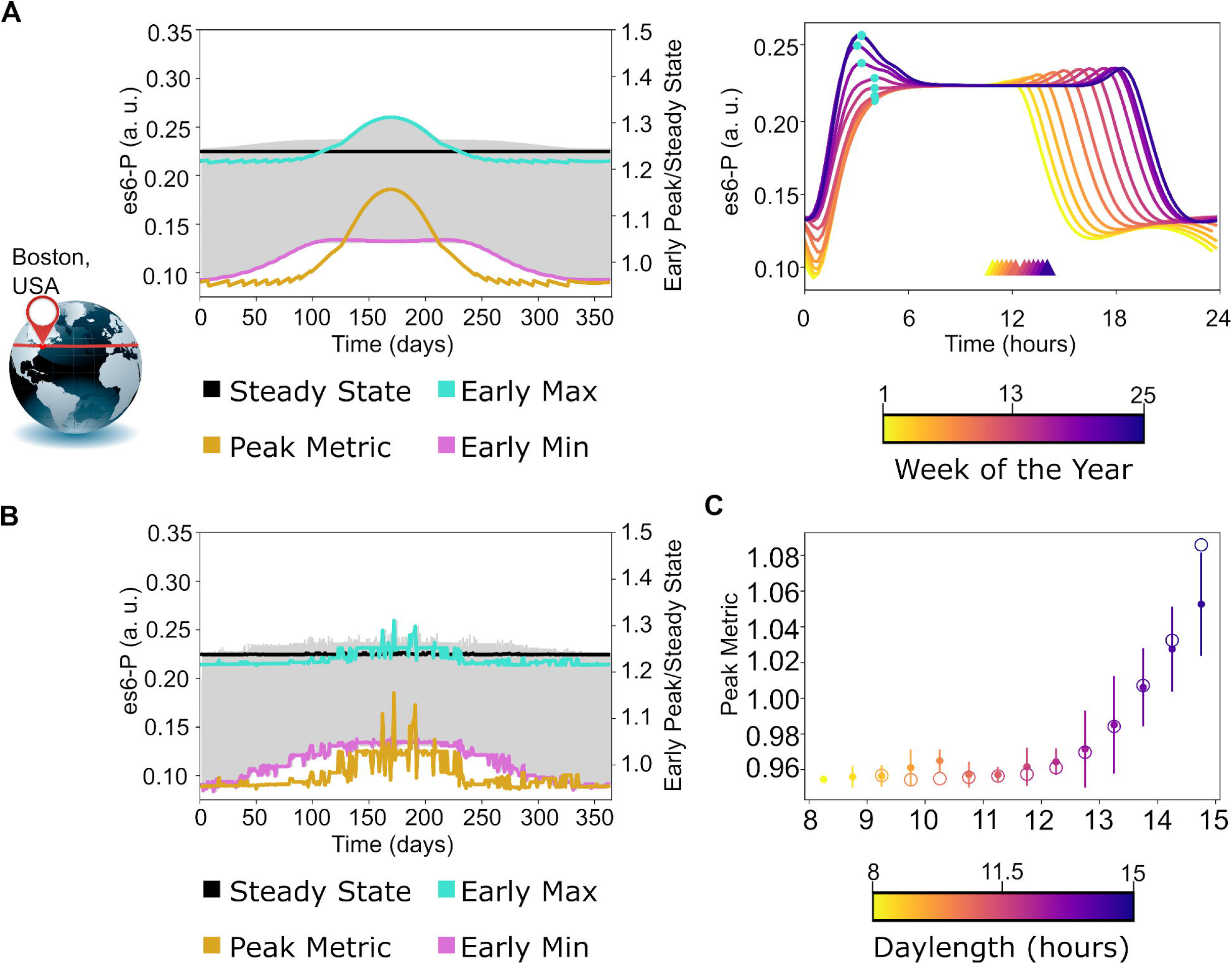
Simulated amounts of eS6-P in response to transient and seasonal changes in daylength over a year. **(A)** Simulation of eS6-P across a full year using daylength data for Boston, Massachusetts (the NOAA Solar Calculator). The left figure shows the eS6-P time course over the full year (grey), overlaid with the early-day maximum (green), early-day minimum (pink), and daily steady state (end-of-day, or dusk) level (black). Peak metric (orange) is defined as the ratio of early-day maximum to the daily steady state level. The right figure shows the eS6-P profiles of individual days selected every two weeks from Day 7 to Day 175 of the year. Time 0 is the actual dawn of each day. The color of each curve corresponds to the week number and is correlated with the daylength from short (yellow) to long (purple). Triangles indicate the time at which the end-of-day eS6-P level was measured, and blue dots indicate the position of the early-day peak. **(B)** Simulation of eS6-P over a year using the full year daylength data derived from Harvard Forest radiometry data [26]. The eS6-P time course over the full year (grey) is overlaid with the early-day maximum (green), early-day minimum (pink), and daily steady state (end-of-day, or dusk) level (black). Peak metric (orange) is defined as the ratio of early-day maximum to the daily steady state level. **(C)** The average peak metric (ratio of the early day maximum of eS6-P to the eS6-P levels at the dusk) for NOAA (open circle) and Harvard Forest (closed circle) days binned according to their realistic daylength (every half hour from 8 to 15 hours). Error bars indicate the standard deviation of Harvard Forest days in each bin. The color of the points and bars is correlated with daylength from short (yellow) to long (purple). The globe on the left shows the latitude of the position that the daylength was simulated for.

The sensitivity of early eS6-P responses to daylength variations raises a question whether this detection of daylength variation is robust with respect to fluctuations of light conditions due to, for example, temporary shading by taller plants, and well as fluctuations in molecular concentrations. We therefore introduced high frequency white noise to the variables describing either the amount of transmitted light or eS6-P itself. We used mutual information to quantify the signal transmitted from varied daylength to the early eS6-P response. We found that more than 50% of the mutual information (compared to the noise-free condition with a finite number of bins) was retained (>1 bit) in the presence of significant fluctuations of light or eS6-P (amplitude parameter *μ* = 5, or about 23% coefficient of variation in light signal) (Fig. S6). This result suggests that the system is robust with respect to the rapid fluctuations of light or molecular concentrations, while it maintains its capacity to detect daylength variations. Intuitively, the peak of eS6-P in long days is driven by a clock-based, slowly increasing trajectory starting from night, and this slow dynamics serves as a signal integration, or averaging method, to reduce the effect of high frequency noise.

### Dynamical features of the eS6-P circuit represent expression profiles of a large number of genes

Because many genes in *A. thaliana* are co-regulated by the circadian clock and light-dark cycles, we hypothesized that the dynamic features of eS6-P can be observed in the expression patterns of other genes. To identify genes whose expression may resemble the profile of eS6-P, we reanalyzed an *A. thaliana* cyclic gene expression data set reported by Dalchau et al. (8), which is based on previously published microarray data with expression patterns of *A. thaliana* genes under LD and LL conditions (6, 31, 32). By calculating the phase shift of the peak expression between measurements under LD and LL conditions, we identified 126 genes where peak expression appears in the early day under LD (0-6 hours after dawn) and regressed into the night under LL condition (18-24 hours after dawn), emulating the clock driven sensitivity that we observed in our eS6-P model and the underlying experimental data (see Methods). We next reanalyzed our previously published RNA sequencing data for *A. thaliana* genes under clock-deficient condition (CCA1 overexpression) (10), and further refined the list of 126 genes by selecting those that have peak expression during the day and have significantly higher expression during the day than during the night under the clock-deficient condition (see Methods). With these selection criteria based on the two data sets (Dalchau et al. and Missra et al. (8, 10)), we have identified 92 genes of which time course expression profiles are similar to that of eS6-P (Table S3). As expected, genes identified in this manner exhibited expression patterns qualitatively similar to eS6-P dynamics (Fig. 7).

**Fig. 7.**
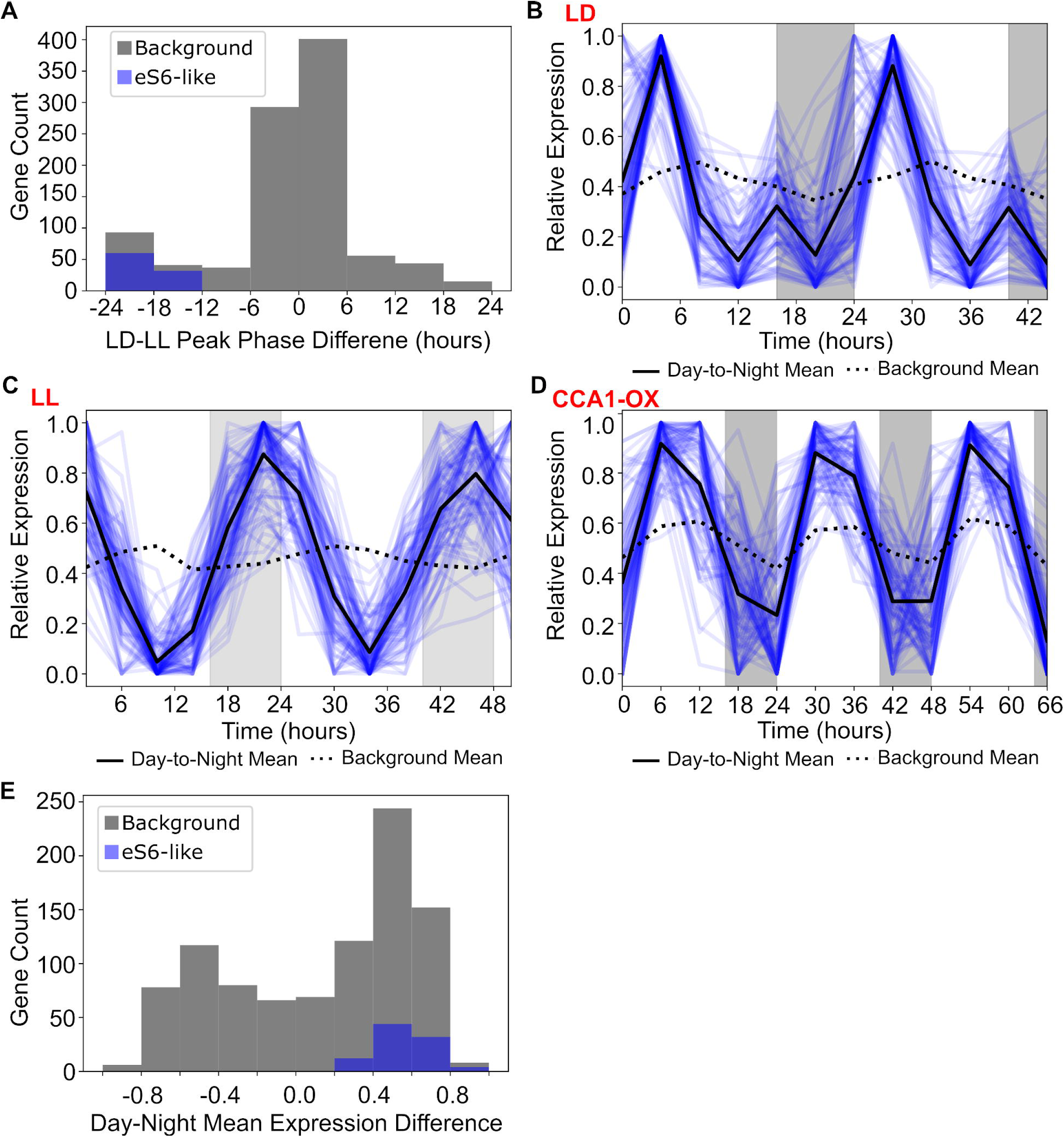
Genes in *A. thaliana* with eS6-P like expression patterns. **(A)** The distribution of the peak phase shift between LD and LL conditions for 92 genes for which phases of expression profiles are similar to those of eS6-P (blue). Equivalent phase shifts for all cyclic genes in the list generated by Dalchau et al. (8) are in gray. **(B-D)** The mRNA expression patterns of 92 genes with phases of expression profiles similar to those of eS6-P for wildtype under LD (B), wildtype under LL (C) and CCA1-overexpression under LD (D) conditions. Time course expression profiles of individual genes are shown as colored curves while the solid black line shows the average expression of all 92 genes. The dotted black line is the average expression pattern of all cyclic genes in the list generated by Dalchau et al. (8). Expression values in all panels were normalized from 0 to 1, such that 0 corresponds to the minimum and 1 correspond to the maximum for each individual gene. **(E)** Distributions of the difference in average mRNA expression level between day and night of 92 eS6-P like genes (blue) and all cyclic genes in the list generated by Dalchau et al. (8) (gray) under CCA1 overexpression condition. Positive values indicate greater average expression during the day and negative values indicate greater average expression during the night.

Dalchau et al. (8) classified genes with cyclic expression patterns into three categories based on whether the clock or the light-dark cycle dominates the amplitude of the oscillation in mRNA levels: clock-dominant, light-dominant and co-regulated. Note that all three types of genes can be influenced by both the clock and the light-dark cycles, and the category ‘co-regulated’ here has a more stringent definition in terms of the amplitudes (see Methods). Not surprisingly, the time course profile of eS6-P is categorized as ‘co-regulated’ based on the definition of Dalchau et al. (see Methods). We found that out of the 92 eS6-P like genes, 66 are classified as ‘clock-dominant’, while 25 are ‘co-regulated’ and only 1 is ‘light-dominant’, suggesting the clock plays a prime role in generating this pattern of expression.

We next performed gene ontology (GO) enrichment analysis with the 92 eS6-P like genes using the set of all annotated genes in *A. thaliana* as a background. We found that eS6-P like genes genes are enriched for light response, photosynthetic regulation, and chloroplast localization, as expected from their responsiveness to light (Table S4). Many of the same biological process and cellular compartment terms are also enriched amongst the set of all genes that show cyclic expression patterns (Table S5). However, the enrichment of specific molecular functions regarding memebrane transport (zinc, ferrous iron, and protons) and activities (ATPpase and thioredoxin-disulfide reductase) is unique to eS6-like genes, suggesting they represent a more specialized subset of the light-clock regulated genes. Overall, this suggests that eS6-P like behavior, a photoperiod sensitive shift in peak activity around dawn, is associated with specific light-dependent metabolic functions.

These results suggest that the dynamics of eS6-P may represent a broad range of transcriptional, post-transcriptional and post-translational activities in *A. thaliana*, particularly in association with light driven and responsive metabolic functions. While eS6-P is co-regulated by both the light and clock, the phase variation in our model is driven by the clock, and this is consistent with the observation that this pattern is associated with many clock-dominant genes. The similarity of dynamical features among these cellular activities does not indicate a causal relationship, but it raises the possibility that the daylength detection and anticipation features of such dynamics may be used by a large system of molecules in *A. thaliana*. This suggests that the cyclic phosphorylation of eS6 may serve as a key signaling factor or an effector integrating ribosomes into this detection and anticipation system.

## Discussion

### Robustness and sensitivity of the clock with respect to light conditions

In this study, we built a mathematical model that describes the dynamics of eS6-P, a ribosomal post-translational modification that occurs in many eukaryotic species. A feedforward regulatory loop connects light signals to eS6-P through a clock dependent pathway and a clock independent pathway. We found that the response of eS6-P is sensitive to the changes of the light-dark cycles, e.g. the daylength variations that are reflected in the time of the dawn. The circuit therefore enables cells to detect and anticipate such variations which may result from changes in season and/or transient changes in weather. We showed that the circadian clock plays important roles in this information processing function. This feature is different from the well-known function of the clock, which ensures robust anticipation of light-dark cycles even under fluctuating light conditions (33, 34). Our study shows that the clock can be used to robustly detect the variations of light conditions at the beginning of a day, which is a function in addition to its traditional role in generating stable rhythmic cues that counteract environmental fluctuations. The circuit achieves this because light and clock signals synergize with each other only at the early phase of the light-dark cycles. This property of the circuit adds to the remarkably diverse ways that the clock may be used by organisms. Furthermore, it might be a fitness advantage for plants to stabilize one group of molecular activities with the clock, while making other activities sensitive to the light conditions using the clock as a reference.

### Physiological functions of external coincidence

Although the sensitivity of eS6-P to light around dawn has not been investigated comprehensively, experimental data showed that eS6-P rises quickly in response to light (18, 22), as anticipated by our model. The general dynamical feature of eS6-P as a result of clock-light signal integration is consistent with other known signaling events in which the coincidence of internal, clock-derived signals and external light signals drives prominent peaks of molecular activities in plants (8, 9, 35). For example, according to the classic ‘external coincidence’ mechanism for photoperiodic (seasonal) flowering, the clock drives up the expression of the regulator of flowering, CONSTANS, late in the day. In long-day plants, if CO happens to be exposed to light late in the day, CO is activated and triggers flowering, whereas if CO is met by darkness, as is the case in the winter, flowering remains suppressed (9). Thus, the external coincidence model allows plants to detect variation in photoperiod at the end of the day. Our work suggests another implementation of external coincidence, where variation in light conditions is detected at the beginning of the day by sensing photoperiod sensitive shifts in the phase of circadian genes(26, 28). For CO at the end of the day, coincidence of clock and light signals activates CO while darkness is incoherent with the clock, and represses CO. For comparison, according to our model eS6-P is most sensitive to changes in light conditions at the beginning of the light period because the negative effect of the clock and the potentially positive effect of light are incoherent. We propose that eS6-P helps the plant to sense variation in the onset of light, as a result of cloud cover or shading by other objects in the morning, and adjust its physiology accordingly.

### Versatile performance objectives of feedforward loops

Previous studies have demonstrated multiple functions of feedforward loops in terms of systems dynamics, including accelerating responses, fold-change detection, adaptation to constant signals and filtering noise (36-39). In addition, it was shown that a feedforward loop can perform a ‘counting’ function that transforms oscillatory input to stable signals (40). Our study shows a previously underappreciated function of a particular type of feedforward loop containing an intrinsic autonomous oscillator. This system not only detects the phase and period variations of the oscillatory inputs by combining the signals from the external and internal oscillations, but also memorizes the altered response in the following cycles, which may serve to anticipate additional perturbations. The latter feature may be useful for plants to predict weather through encoding the information in the light conditions of the previous day. This result further demonstrates the versatility of the functions of the feedforward loop. Future work is needed to determine whether this phase-detection mechanism can be integrated with other known functions of feedforward loops.

### Anticipation of changes in environmental conditions

Phosphorylation of eS6 occurs in all organisms where it has been examined, including yeast, plants, and humans (12-14). A previous study showed dramatically increased translation activity in mammalian cells with a phosphorylation-deficient eS6, suggesting a possible role of eS6-P in controlling general resource allocation in cells (15). If this potential function of eS6-P were conserved across kingdoms, then our work would further suggest that eS6-P might tune some aspect of protein synthesis in response to variable light condition. This function, while still to be demonstrated in plants, would not be far-fetched given that translation requires a substantial input in cellular energy, which may be depleted at the end of night. This detection and anticipation of daylength may help plants to prepare for days with particular patterns of light exposures. It has been shown that growing yeast cells can allocate resources to anticipate favorable or unfavorable environmental conditions, and the choice of these anticipations may be made with rhythmic dynamics (41, 42). Regardless of the specific cellular functions of eS6-P, our work sheds light on the rich dynamics of eS6-P under fluctuating environmental conditions. Moreover, the remarkable information processing characteristic of the signaling circuit that controls eS6-P has the capacity to detect and memorize critical cues to allow plants to adapt to a dynamical light environment.

## Methods

### Framework of mathematical models

We modeled gene regulatory networks that control eS6-P using ordinary differential equations (ODEs). The network topologies are shown in Fig. 1. Because most interactions involving high-order molecular interactions, including transcriptional regulation and multisite phosphorylation, we used a generic form of nonlinear ODEs suitable for describing both gene expression and molecular interaction networks (43-46). Each ODE system in the model has the form:

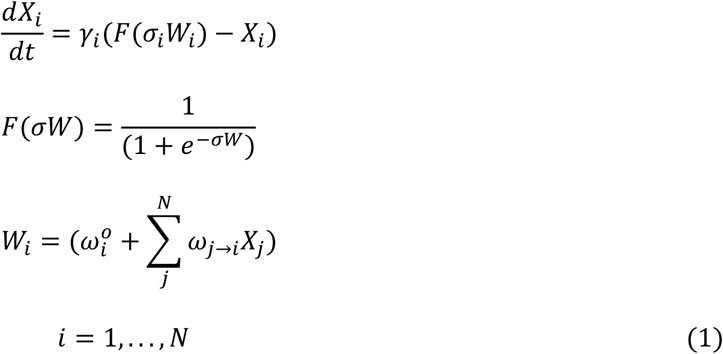

Here, *X*_*i*_ is the activity of protein *i*. On a time scale 1/*γ*_*i*_, *X*_*i*_(*t*) relaxes toward a value determined by the sigmoidal function, *F*, which has a steepness set by *σ*. The basal value of *F*, in the absence of any influencing factors, is determined by 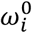. The coefficients *ω*_*j*→*i*_ determine the influence of protein *j* on protein *i. N* is the total number of proteins in the network. Activity values are scaled from 0 (absent) to 1 (saturation). For eS6-P, we modeled a single site phosphorylation (S237) that was assumed to be independent of other phosphorylation sites, so we used first order reaction rate law that describes the phosphorylation and dephosphorylation. All variables and parameters are dimensionless. One time unit in the simulations corresponds to approximately 2.53 hours (9.5 unit per cycle), but we transformed the time unit to 1 hour for all subsequent analyses and visualizations.

The clock component of our model describes four species (LHY/CCA1, EC, PRR9/7, and PRR5/1), and the light-sensing pathway has two species (TOR and S6K). Together with eS6-P, our full Clock+Light model has seven ODEs in total. Numerical solutions to the ODEs were obtained with Tellurium (47). Computer code for simulating the models under various conditions and reproducing key figures is available at: https://github.com/panchyni/eS6_Models.

### Parameter estimation

Because the clock component of the model is independent of other elements except for the light input, we first fit the clock components of the model (4 ODEs) by applying qualitative criteria to ensure that it properly replicated features of the circadian clock [18]. We approached optimization the same way as in Locke at al. (5) and De Caluwe et al. (48) by using objective functions which impose costs on deviating from expected, qualitative behavior. Specifically, we penalized the model for (1) having the LHY/CCA1 component peak more than 1 hour before or after dawn in long (16L:8D) days, (2) having the same deviation described in (1) in 12L:12D days, (3) having a period under constant light outside of 24 to 25 hours, (4) having a period under constant darkness outside of 24 to 28 hours, and (5) having an amplitude of less than 0.1 (i.e. 10% of maximum activity) in any component. These criteria defined by intervals of desired output were used to construct an objective function to evaluate the models. When calculating the objective score, each of the five terms is zero if the output falls in the interval and is the squared difference between the output and the boundary of the interval if the output falls outside the interval. Before optimizing the model parameters, we defined a hyperbox in the parameter space that is bounded by biologically plausible parameter ranges, particularly with regard to the direction of regulation and balance of rates. A population of 40 vectors, each containing 24 parameters, generated by Latin hypercube sampling (LHS), were used as the initial estimation. Starting with this population, we implemented Differential Evolution (DE) optimization algorithm (49, 50) with a mutation rate between 0.35 and 0.65 and a crossover rate of between 0.75 and 0.95, and let the optimization run for 15000 generations of DE or the convergence of the evolution. Similar to the approach used by De Caluwe et al. (48), we then compared each component to expression data from Diurnal Database (27) which were normalized onto a 0 to 1 scale to ensure the estimated component activity approximated the proper phase and shape of actual clock components (Fig. S2). We performed 250 runs of such optimization to ensure that the performance of the selected parameter set can be reproduced. For fitting eS6-P, we selected among viable models which accurately reflected clock behavior and selected the one with the greatest dynamic range of among the clock components.

Next, we used the time course measurement of eS6-P to estimate the parameters controlling the regulation of eS6-P by clock components (LHY/CCA1, EC, PRR97, and PRR51 which correspond to C1, C2, C3, and C4 respectively) and the light induced TOR pathway (18). The data set has Western Blot quantification of eS6-P for 78 hours at a 6-hour interval. In the model, the eS6-P variable describes the percentage of the amount of eS6-P with respect to the total eS6. We therefore inferred the fractions of eS6-P from the Western Blot quantification of the experiment and scaled these values based on an approximation of the eS6-P saturation value inferred from Williams et al. (51). 2D-gel electrophoresis data from this study suggest that a majority, but not all eS6 undergoes phosphorylation and that phosphorylation spreads across several sites which may or may not be phosphorylated at the same time (see Fig. 4 and Table 1 in (51)). As the data were not quantified in the original study and estimations based on image analysis were broad (15-77% per site), we chose a conservative estimate of 0.40 as the saturating value. This normalization gives the raw data a range of 0.01 to 0.4 and pooled data (described below) a range of average values from 0.06 to 0.31. This procedure allows us to avoid unrealistic assumptions with either no phosphorylated (eS6-P = 0) or fully phosphorylated (eS6-P = 1) in the system. Because these values are approximate and mainly meant to avoid fitting to unrealistic, extreme values of eS6-P, the model describes eS6-P in arbitrary units (a. u.) rather than actual percentages.

We regularized the transformed data by pooling the data points from the same Zeitgeber time, and we used the pooled mean and variance for each time point to calculate the log likelihood. Constant light was emulated in the model by fixing the value of light to 1 and the CCA1 overexpression by fixing the basal production of CCA1 to 50. To examine steady state behavior under each condition, we entrained the model with one day of darkness followed by ten days of the given condition, before comparing the model to the experimental data on the twelfth day. The objective function for the parameter optimization was defined by the log-likelihood function that quantifies the fit of the simulation results to the experimental data with variabilities. Log likelihood estimates for all time points of eS6-P contribute to the objective function equally in an additive manner. As with clock components, we used the DE based optimization algorithm to estimate the remaining parameters. All algorithmic parameters were the same as the optimization for the clock component, expect for the maximum number of generations (5000). We performed 1400 optimization runs, and among the optimized parameter sets we selected the best preforming model with a moderate peak to perform subsequent analyses. Note that the majority of the models in the lowest 5th percentile of likelihood values exhibited the same behavior as the chosen model, including cycling under all three experimental conditions and an early eS6-P peak under LD conditions (Fig. S3A). Models with larger likelihood scores (the next 5 percentiles) show neither this earlier peak nor cycling under constant light (Fig. S3B). A full list of model parameters can be found in Table S1.

### Applying a detailed clock to our eS6-P model

In addition to our simplified 4-component clock model, we also tested our eS6-P model using a more detailed circadian clock component from De Caluwe et al. (48), which has the same overall network topology but explicitly considers protein and mRNA concentrations (Fig. S4). We added our equation for eS6-P regulation to this full clock model, and used the set of same optimized parameters that we obtained from the simplified model to simulate eS6-P with the light-dark cycle. The CCA1 constitutive overexpression mutant was simulated by removing the clock and light regulated components from the equations for CCA1/LHY transcription (i.e. setting the production rate to maximum). We used the protein concentration of each pair of clock components, representing the mean protein concentration of the two gene products, in place of our generalized clock component activity values. We scaled the concentration values to a range of [0,1] using the minimum and maximum values of that protein pair. However, in the case of CCA1 overexpression, we assumed that the concentration is effectively saturated at the maximum wild-type value, as the average value of CCA1/LHY and PRR9/7 increased by more than five-fold. All parameters for eS6 regulation were the same as in the model in Fig. 1, except for the regulatory effect of the TOR pathway which was increased by 10% to avoid an unrealistic saturation point of eS6-P under wild-type and CCA1 overexpression conditions.

### Alternative models

To illustrate the unique performance of the Clock+Light model that contains the clock component, we built three alternative models that describe plausible ways in which the eS6-P can transmit the light signal to achieve detection of daylength variations at the beginning of a day. Note that it is trivial for a system to detect daylength variations at the end of a day (e.g. through integration of, or slow response to, light signals), and this detection may not be as useful as the early day detection in terms of anticipating environmental changes. The three alternative models are: 1) a linear circuit that transmits light signals to eS6-P (Fig. 3A, top panel), 2) an incoherent feedforward loop (IFFL) that produces an early day peak similar to what the Clock+Light model does under LD condition (Fig. 3A and B, middle panels), and 3) a coherent feedforward loop (CFFL) that allows slow decline of eS6-P upon the withdrawal of light signals (Fig. 3A, lower panel). Parameter values of these models were manually chosen to give characteristic responses (Fig. 3A, Table S2).

### Mutual information between daylength variations and eS6-P responses

To examine the transmitted information from daylength variations (signal) to eS6-P abundance (response) when the system is subject to external or internal noise, we introduced uncorrelated multiplicative white noise to ODEs for the light and eS6-P as *dx*/*dt* = *f*(*x*_1_, *x*_2_, …, *x*_*n*_) + *μ*_*x*_ · *x* · *dW*, where *dW* is a Wiener process that can be discretized as 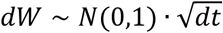 where *N*(0,1) denotes a normally distributed random variable with zero mean and unit variance, and *μ*_*x*_ represents the amplitude of the fluctuation. To quantify the information transmission, we used mutual information between signal *S* and response *R* (52). In the eS6-P system, *S* represents the perturbation of the time (Δt_ND_) at which the system receives light signal from a normal light-dark cycle (e.g. 12hr light and 12hr dark), and *R* represents the eS6-P abundance. The responses were the maximum eS6-P levels in the 4-hour time window after dawn. The calculation of the mutual information was performed with the discretized form (53):

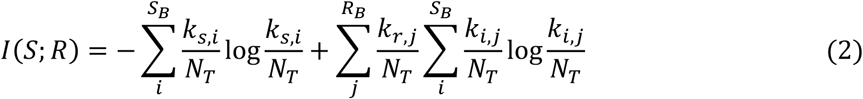

Here, signals values were assigned to *S*_*B*_ bins, and the response values were assigned to *R*_*B*_ bins. By binning all signals and responses, we constructed a contingency matrix of which each entry is the number of observations from the simulated data that correspond to that particular signal-response pair. *N*_*T*_ is the sum over all entries in the table, and *k*_*i,j*_ is the number of instances of signal *i* that resulted in response *j*. In this study, 21 signal bins and 10 response bins were used. Using more response or signal bins will likely increase the mutual information, so our analysis focuses on comparisons of lower bounds of the mutual information. The bounds of the response bins were determined by the minimal and maximal responses in the absence of the noise. For each signal value, 200 simulations were performed.

### Photoperiod data processing

We obtained a copy of the NOAA Solar Calculator (https://www.esrl.noaa.gov/gmd/grad/solcalc/calcdetails.html) and generated approximated day length information (i.e. Sunlight Duration) for a full year for three locations: Oslo, Norway (59.91 Latitude, 110.75 Longitude), Praia, Cape Verde (14.92 Latitude, -23.51 Longitude), and Boston, Massachusetts (42.35 Latitude, -71.05 Longitude). For inclusion in our model, we fit a model of day-night variation to each data set with the following form:

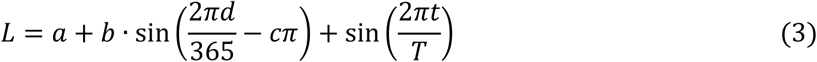

where the rightmost sine term represents a base 12-hr light/dark cycles and the leftmost sine term modulates this average day based on the time of the year. Here, *d* is the day of the year, *t* is the time of day in hours and *T* is the period of the day in hours. The parameters *a, b*, and *c*, are fit such that the fraction of daylight (*L* > 0) conforms to the NOAA estimations for each day (values for each location are listed in Table S6). With this idealized year-long data, the time for analysis of daily response is based the natural light condition rather than any artificially defined time. We also obtained measurement of radiation data in the Harvard Forest (30) including downward oriented photosynthetically active radiation (par.down), which we used as an approximation for daylight. A threshold value (9.0) was used to define day/night using par.down and was chosen such that it minimized the average difference between the inferred hours of daylight and NOAA estimations of day length across all days binned by month. For inclusion in our model, we smoothed radiation data from 2006 (which had the fewest missing values) using cubic splines and applied the threshold to generate binary day/night values.

### Identification of genes with time-series expression profiles similar to eS6-P dynamics

To identify genes that show similar cyclic patterns to eS6-P dynamics under LD, LL and CCA1 overexpression conditions, we first selected genes identified by Dalchau et al. (8) where the peak expression in long-day was during the early day (ZT 0-6 hours) but the peak expression in constant light was during subjective night (ZT 18-24 hours). There were 126 genes that satisfy these conditions. We then used the RNA sequencing data from Missra et al. (measurement under CCA1 overexpression condition) to further filter these genes. We selected genes that had both peak expression during the day, and a difference between average expression during the day and night of at least 25% of the peak value under CCA1 overexpression condition. To visualize the time course profiles of these genes, we used data from Diurnal Database for LD (27), Edwards et al. for LL (32), and Missra et al. for CCA1 overexpression (10).

For gene ontology (GO) analysis, we used all 92 eS6-P like genes and tested for enrichment against all *A. thaliana* genes using PANTHER ((54); available through http://geneontology.org/). The same procedure we used when testing all genes identified by Dalchau et al.(8). *p*-values were calculated using the Fisher’s exact test and multiple test correction was done using Benjamini-Hochberg method with a false discovery rate threshold of 0.05.

Dalchau et al. classified the genes with cyclic expression patterns into three categories: light-dominant, clock dominant and co-regulated (8). To examine which category the eS6-P profile would belong to, we used the same approach to classifying light/clock regulation as Dalchau et al. by comparing the ratio of eS6-P amplitude driven by the clock (constant light) to that driven by the light-dark cycle. Briefly, amplitude difference between LL condition and LD condition was used to determine whether light or clock dominates the regulation of each gene. With our eS6-P data, the amplitude under the LD condition is higher than that under LL condition by 2.17 fold (difference in Bode magnitude = -3.3 decibels, dB), which categorizes eS6 phosphorylation as a process co-regulated by light and clock according to the cutoffs (±7dB difference in Bode magnitude), although light regulation is favored.

## Author Contributions

Designed research: TH. Performed research: NP and TH. Analyzed data and wrote the manuscript: NP, AGV and TH.

## Conflict of Interest

The authors declare no conflict of interest.

## Notes

### Competing Interest Statement

The authors have declared no competing interest.

