## SupplementaryMaterial for "Early detection of daylengths with a feedforward circuit coregulated by circadian and diurnal cycles"

**Table S1. Parameter values of the Clock+Light model**

| Parameter | Description | Values |
|---|---|---|
| γLHY/CCA1 | Timescale of LHY/CCA1 | 0.87 |
| σLHY/CCA1 | Steepness of sigmoidal function of LHY/CCA1 | 2.57 |
| ωLHY/CCA1,0 | Basal production rate of LHY/CCA1 | 8.27 |
| ωLHY/CCA1,PRR9/7 | Activation/inhibition strength of PRR9/7 on LHY/CCA1 | -9.90 |
| ωLHY/CCA1,PRR5/1 | Activation/inhibition strength of PRR5/1 on LHY/CCA1 | -5.05 |
| ωLHY/CCA1,Light | Activation/inhibition strength of Light on LHY/CCA1 | -1.15 |
| γPRR9/7 | Timescale of PRR9/7 | 0.22 |
| σPRR9/7 | Steepness of sigmoidal function of PRR9/7 | 6.06 |
| ωPRR9/7,0 | Basal production rate of PRR9/7 | 1.58 |
| ωPRR,LHY/CCA1 | Activation/inhibition strength of LHY/CCA1 on PRR9/7 | 4.47 |
| ωPRR9/7,PRR5/1 | Activation/inhibition strength of PRR5/1 on PRR9/7 | -1.56 |
| ωPRR9/7,EC | Activation/inhibition strength of EC on PRR9/7 | -9.88 |
| ωPRR9/7,Light | Activation/inhibition strength of Light on PRR9/7 | 1.53 |
| γPR5/1 | Timescale of PRR5/1 | 3.74 |
| σPRR5/1 | Steepness of sigmoidal function of PRR5/1 | 9.86 |
| ωPRR9/7,0 | Basal production rate of PRR5/1 | 7.35 |
| ωPRR5/1,LHY/CCA1 | Activation/inhibition strength of LHY/CCA1 on PRR5/1 | -10.00 |
| ωPRR5/1,PRR5/1 | Activation/inhibition strength of PRR5/1 on PRR5/1 | -7.59 |
| γEC | Timescale of EC | 9.79 |
| σEC | Steepness of sigmoidal function of EC | 4.06 |
| ωEC,0 | Basal production rate of EC | 7.12 |
| ωEC,LHY/CCA1 | Activation/inhibition strength of LHY/CCA1 on EC | -9.68 |
| ωEC,PRR5/1 | Activation/inhibition strength of PRR5/1 on EC | -5.08 |
| ωEC,EC | Activation/inhibition strength of EC on EC | -0.73 |
| γSK6 | Timescale of SK6 | 2.54 |
| σSK6 | Steepness of sigmoidal function of SK6 | 4.43 |
| ωSK6,0 | Basal production rate of SK6 | -1.23 |
| ωSK6,TOR | Activation/inhibition strength of TOR on SK6 | 3.89 |
| γTOR | Timescale of TOR | 1.11 |
| σTOR | Steepness of sigmoidal function of TOR | 1.00 |
| ωTOR,0 | Basal production rate of TOR | -2.62 |
| ωTOR,Light | Activation/inhibition strength of Light on TOR | 7.70 |
| γRPS6 | Timescale of eS6 | 5.26 |
| σRPS6_phos | Steepness of sigmoidal function of eS6 phosphorylation | 1.09 |
| σRPS6_dephos | Steepness of sigmoidal function of eS6 dephosphorylation | 1.55 |
| ωRPS6,0_phos | Basal production rate of eS6 phosphorylation | -2.28 |
| ωRPS6,0_dephos | Basal production rate of eS6 dephosphorylation | 6.52 |
| ωRPS6,SK6 | Activation/inhibition strength of SK6 on eS6 phosphorylation | 1.45 |

| | | |
|---|---|---|
| ωRPS6,LHY/CCA1 | Activation/inhibition strength of LHY/CCA1 on eS6 dephosphorylation | -8.63 |
| ωRPS6,PRR9/7 | Activation/inhibition strength of PRR9/7 on eS6 dephosphorylation | 2.94 |
| ωRPS6,PRR5/1 | Activation/inhibition strength of PRR5/1 on eS6 dephosphorylation | 6.55 |
| ωRPS6,EC | Activation/inhibition strength of EC on eS6 dephosphorylation | -8.58 |

**Table S2. Parameter values of alternative models**

| Parameter | Description | Linear Circuit Model | IFFL Model | CFFL Model |
|---|---|---|---|---|
| $\gamma_A$ | Timescale of A | 1 | 1 | NA |
| $\sigma_A$ | Steepness of sigmoidal function of A | 1 | 5 | NA |
| $\omega_{A,0}$ | Basal production rate of A | -1.9 | -1 | NA |
| $\omega_{A,Light}$ | Activation/inhibition strength of light on A | 3 | 3 | NA |
| $\gamma_B$ | Timescale of B | NA | 1 | 0.3 |
| $\sigma_B$ | Steepness of sigmoidal function of B | NA | 10 | 5 |
| $\omega_{B,0}$ | Basal production rate of B | NA | -1 | -1.9 |
| $\omega_{B,Light}$ | Activation/inhibition strength of light on B | NA | 3 | 3 |
| $\gamma_C$ | Timescale of C | NA | 2 | 0.3 |
| $\sigma_C$ | Steepness of sigmoidal function of C | NA | 10 | 1 |
| $\omega_{C,0}$ | Basal production rate of C | NA | -0.6 | -1 |
| $\omega_{C,Light}$ | Activation/inhibition strength of | NA | NA | 3 |
| $\omega_{C,B}$ | Activation/inhibition strength of | NA | 1 | NA |
| $\gamma_{RPS6}$ | Timescale of eS6 | 10 | 10 | 10 |
| $\sigma_{RPS6\_phos}$ | Steepness of sigmoidal function of eS6 phosphorylation | 1 | 10 | 1 |
| $\sigma_{RPS6\_dephos}$ | Steepness of sigmoidal function of eS6 dephosphorylation | 1 | 1 | 1 |
| $\omega_{RPS6,0\_phos}$ | Basal rate of eS6 phosphorylation | -6.3 | -5 | -6.3 |
| $\omega_{RPS6,0\_dephos}$ | Basal rate of eS6 dephosphorylation | 2 | 2 | 2 |
| $\omega_{RPS6\_phos,A}$ | Activation/inhibition strength of A on eS6 phosphorylation | 10 | 10 | NA |
| $\omega_{RPS6\_dephos,A}$ | Activation/inhibition strength of A on eS6 dephosphorylation | 0 | 0 | NA |
| $\omega_{RPS6\_phos,B}$ | Activation/inhibition strength of B on eS6 phosphorylation | NA | NA | 10 |
| $\omega_{RPS6\_dephos,B}$ | Activation/inhibition strength of B on eS6 dephosphorylation | NA | NA | 0 |
| $\omega_{RPS6\_phos,C}$ | Activation/inhibition strength of C on eS6 phosphorylation | NA | -6.8 | 10 |
| $\omega_{RPS6\_dephos,C}$ | Activation/inhibition strength of C on eS6 dephosphorylation | NA | 0 | 0 |

**Table S3. Dalchau Genes with eS6-P expression in LD, LL, and CCA1 conditions**

| Locus number | Difference in Bode magnitude | SD phase | LD phase | LL phase | Dalchau Category |
|---|---|---|---|---|---|
| AT3G24190 | 1.9812 | 3 | 4 | 20.23 | Co-regulated |
| AT2G31790 | -4.8845 | 5 | 4 | 20.23 | Co-regulated |
| AT4G04610 | -5.102 | 4 | 5 | 20.43 | Co-regulated |
| AT1G05690 | 3.9141 | 0 | 2 | 20.59 | Co-regulated |
| AT5G19850 | 4.8354 | 4 | 4 | 20.97 | Co-regulated |
| AT1G33110 | 5.606 | 1 | 1 | 21.33 | Co-regulated |
| AT1G50020 | 1.0719 | 5 | 5 | 21.59 | Co-regulated |
| AT4G17840 | 5.8182 | 3 | 5 | 21.6 | Co-regulated |
| AT4G26860 | 5.445 | 23 | 0 | 21.78 | Co-regulated |
| AT2G37240 | 5.5772 | 5 | 5 | 21.79 | Co-regulated |
| AT4G34720 | 0.9715 | 5 | 5 | 21.87 | Co-regulated |
| AT5G15450 | 4.9274 | 4 | 4 | 21.93 | Co-regulated |
| AT5G40390 | 3.0197 | 5 | 5 | 22.01 | Co-regulated |
| AT1G65490 | 6.0176 | 0 | 1 | 22.08 | Co-regulated |
| AT1G06430 | 5.4129 | 5 | 5 | 22.08 | Co-regulated |
| AT2G04350 | 2.706 | 3 | 5 | 22.2 | Co-regulated |
| AT1G10370 | 4.7385 | 5 | 4 | 22.66 | Co-regulated |
| AT4G08870 | 3.3578 | 5 | 6 | 22.87 | Co-regulated |
| AT4G37760 | 6.71 | 5 | 5 | 23.15 | Co-regulated |
| AT2G41290 | 6.7407 | 4 | 4 | 23.36 | Co-regulated |
| AT5G13770 | 5.9211 | 0 | 2 | 23.44 | Co-regulated |
| AT1G18060 | 3.153 | 5 | 5 | 23.48 | Co-regulated |
| AT1G29700 | -1.22 | 5 | 5 | 23.52 | Co-regulated |
| AT2G36870 | 1.7935 | 5 | 5 | 23.59 | Co-regulated |
| AT2G22240 | -0.0301 | 4 | 5 | 23.94 | Co-regulated |
| AT1G10850 | 7.882 | 0 | 2 | 19.6 | Clock Dominant |
| AT1G19970 | 15.678 | 3 | 4 | 20.4 | Clock Dominant |
| AT3G21390 | 18.8453 | 1 | 3 | 20.6 | Clock Dominant |
| AT1G04550 | 10.1137 | 22 | 0 | 21.17 | Clock Dominant |
| AT4G28290 | 25.0479 | 3 | 3 | 21.31 | Clock Dominant |
| AT1G15060 | 15.2501 | 3 | 5 | 21.41 | Clock Dominant |
| AT2G33250 | 20.1534 | 1 | 1 | 21.47 | Clock Dominant |
| AT5G62220 | 15.1542 | 1 | 2 | 21.5 | Clock Dominant |
| AT1G70610 | 9.0462 | 5 | 5 | 21.57 | Clock Dominant |
| AT2G46735 | 25.1114 | 4 | 4 | 21.67 | Clock Dominant |
| AT1G50250 | 23.688 | 4 | 5 | 21.7 | Clock Dominant |
| AT2G04039 | 17.5514 | 3 | 3 | 21.77 | Clock Dominant |
| AT1G79520 | 9.0837 | 3 | 5 | 21.77 | Clock Dominant |

| | | | | | |
|---|---|---|---|---|---|
| AT4G14240 | 8.9341 | 5 | 6 | 21.86 | Clock Dominant |
| AT2G15020 | 14.1699 | 4 | 4 | 21.9 | Clock Dominant |
| AT3G27170 | 19.4057 | 2 | 3 | 21.95 | Clock Dominant |
| AT4G32770 | 17.2847 | 4 | 5 | 22.06 | Clock Dominant |
| AT5G03555 | 8.6669 | 23 | 1 | 22.14 | Clock Dominant |
| AT3G01180 | 12.6212 | 4 | 5 | 22.17 | Clock Dominant |
| AT1G19450 | 19.8414 | 3 | 1 | 22.21 | Clock Dominant |
| AT3G27050 | 40.3256 | 1 | 3 | 22.3 | Clock Dominant |
| AT2G46450 | 15.7447 | 3 | 1 | 22.32 | Clock Dominant |
| AT4G17530 | 12.2573 | 0 | 2 | 22.37 | Clock Dominant |
| AT4G24700 | 10.8052 | 1 | 2 | 22.37 | Clock Dominant |
| AT4G32190 | 18.553 | 3 | 2 | 22.4 | Clock Dominant |
| AT1G70985 | 14.1352 | 5 | 5 | 22.43 | Clock Dominant |
| AT2G34460 | 7.0858 | 3 | 5 | 22.43 | Clock Dominant |
| AT3G23920 | 15.9258 | 3 | 3 | 22.49 | Clock Dominant |
| AT2G04570 | 17.8394 | 2 | 2 | 22.56 | Clock Dominant |
| AT2G39450 | 11.2944 | 3 | 3 | 22.6 | Clock Dominant |
| AT3G01550 | 31.8332 | 3 | 5 | 22.63 | Clock Dominant |
| AT5G48930 | 14.2 | 1 | 2 | 22.69 | Clock Dominant |
| AT3G44100 | 8.0675 | 5 | 5 | 22.74 | Clock Dominant |
| AT1G66330 | 21.1295 | 3 | 5 | 22.74 | Clock Dominant |
| AT1G07180 | 19.8284 | 3 | 3 | 22.8 | Clock Dominant |
| AT1G64860 | 9.7185 | 2 | 2 | 22.8 | Clock Dominant |
| AT1G01520 | 20.4053 | 4 | 4 | 22.83 | Clock Dominant |
| AT3G23000 | 19.8077 | 5 | 5 | 22.85 | Clock Dominant |
| AT5G23730 | 73.6304 | 3 | 3 | 22.86 | Clock Dominant |
| AT2G26690 | 19.6111 | 1 | 3 | 22.87 | Clock Dominant |
| AT2G16070 | 42.1697 | 3 | 3 | 22.93 | Clock Dominant |
| AT3G17800 | 11.6961 | 3 | 5 | 22.96 | Clock Dominant |
| AT1G04530 | 16.9199 | 3 | 3 | 23.01 | Clock Dominant |
| AT5G58870 | 19.2999 | 3 | 3 | 23.03 | Clock Dominant |
| AT2G24540 | 16.7208 | 3 | 3 | 23.03 | Clock Dominant |
| AT1G22850 | 17.703 | 5 | 5 | 23.04 | Clock Dominant |
| AT4G02420 | 10.873 | 1 | 2 | 23.06 | Clock Dominant |
| AT5G23060 | 16.7202 | 1 | 2 | 23.1 | Clock Dominant |
| AT1G44000 | 15.9371 | 2 | 1 | 23.1 | Clock Dominant |
| AT1G32900 | 35.4202 | 3 | 3 | 23.12 | Clock Dominant |
| AT5G02120 | 28.774 | 1 | 3 | 23.16 | Clock Dominant |
| AT5G50100 | 15.6106 | 5 | 5 | 23.22 | Clock Dominant |
| AT3G61890 | 12.0026 | 1 | 3 | 23.24 | Clock Dominant |
| AT1G18810 | 12.1558 | 4 | 2 | 23.31 | Clock Dominant |
| AT5G52570 | 16.2838 | 2 | 3 | 23.32 | Clock Dominant |
| AT3G11670 | 10.1228 | 2 | 3 | 23.42 | Clock Dominant |

| | | | | | |
|---|---|---|---|---|---|
| AT4G11570 | 15.6497 | 5 | 5 | 23.52 | Clock Dominant |
| AT3G54660 | 22.1698 | 2 | 5 | 23.65 | Clock Dominant |
| AT4G35250 | 16.6058 | 4 | 5 | 23.74 | Clock Dominant |
| AT1G04250 | 10.6132 | 1 | 3 | 23.74 | Clock Dominant |
| AT2G21320 | 17.1625 | 4 | 3 | 23.8 | Clock Dominant |
| AT3G10420 | 30.4161 | 2 | 3 | 23.8 | Clock Dominant |
| AT1G55960 | 22.2348 | 3 | 3 | 23.93 | Clock Dominant |
| AT5G12470 | 29.9632 | 2 | 3 | 23.94 | Clock Dominant |
| AT3G24170 | 13.6387 | 2 | 3 | 23.94 | Clock Dominant |
| AT4G04850 | 14.8517 | 5 | 5 | 23.96 | Clock Dominant |
| AT5G03650 | -12.43 | 5 | 5 | 24 | Light Dominant |

**Table S4. GO Term enriched in eS6-P like genes**

| GO Term | Category | # of Genes | Expected # of Genes | Fold Enrichment | p-value | FDR |
|---|---|---|---|---|---|---|
| starch biosynthetic process | biological process | 4 | 0.11 | 36.12 | 7.11E-06 | 8.52E-03 |
| glucan biosynthetic process | biological process | 5 | 0.38 | 13.19 | 4.87E-05 | 2.43E-02 |
| carbohydrate biosynthetic process | biological process | 7 | 1.03 | 6.82 | 8.85E-05 | 3.53E-02 |
| cellular polysaccharide metabolic process | biological process | 7 | 0.95 | 7.35 | 5.61E-05 | 2.58E-02 |
| cellular carbohydrate metabolic process | biological process | 9 | 1.36 | 6.61 | 1.07E-05 | 7.11E-03 |
| cellular carbohydrate biosynthetic process | biological process | 7 | 0.69 | 10.08 | 7.82E-06 | 7.81E-03 |
| cellular glucan metabolic process | biological process | 7 | 0.71 | 9.84 | 9.09E-06 | 7.78E-03 |
| glucan metabolic process | biological process | 7 | 0.74 | 9.48 | 1.15E-05 | 6.88E-03 |
| starch metabolic process | biological process | 5 | 0.2 | 24.83 | 2.64E-06 | 3.96E-03 |
| regulation of photosynthesis, light reaction | biological process | 3 | 0.09 | 34.38 | 1.24E-04 | 4.13E-02 |
| regulation of photosynthesis | biological process | 5 | 0.14 | 34.65 | 5.73E-07 | 1.14E-03 |
| response to high light intensity | biological process | 4 | 0.21 | 18.92 | 7.64E-05 | 3.27E-02 |
| response to light intensity | biological process | 5 | 0.49 | 10.21 | 1.56E-04 | 4.92E-02 |
| response to light stimulus | biological process | 14 | 2.36 | 5.94 | 1.12E-07 | 6.69E-04 |
| response to radiation | biological process | 14 | 2.44 | 5.74 | 1.70E-07 | 5.08E-04 |
| response to abiotic stimulus | biological process | 20 | 6.96 | 2.87 | 1.59E-05 | 8.66E-03 |
| cellular response to light stimulus | biological process | 5 | 0.44 | 11.29 | 9.89E-05 | 3.70E-02 |
| cellular response to radiation | biological process | 5 | 0.46 | 10.8 | 1.21E-04 | 4.26E-02 |
| chloroplast organization | biological process | 6 | 0.79 | 7.64 | 1.58E-04 | 4.73E-02 |
| plastid organization | biological process | 8 | 1.02 | 7.87 | 9.97E-06 | 7.47E-03 |
| glycogen (starch) synthase activity | molecular function | 2 | 0.01 | > 100 | 1.65E-04 | 3.46E-02 |

| | | | | | | |
|---|---|---|---|---|---|---|
| transferase activity, transferring hexosyl groups | molecular function | 8 | 1.41 | 5.66 | 9.73E-05 | 2.19E-02 |
| ferrous iron transmembrane transporter activity | molecular function | 2 | 0.02 | > 100 | 2.30E-04 | 4.53E-02 |
| inorganic cation transmembrane transporter activity | molecular function | 10 | 1.47 | 6.79 | 2.66E-06 | 1.67E-03 |
| inorganic molecular entity transmembrane transporter activity | molecular function | 11 | 2.43 | 4.53 | 3.46E-05 | 1.09E-02 |
| transmembrane transporter activity | molecular function | 15 | 4.03 | 3.72 | 1.18E-05 | 4.66E-03 |
| transporter activity | molecular function | 15 | 4.26 | 3.53 | 2.21E-05 | 7.76E-03 |
| cation transmembrane transporter activity | molecular function | 11 | 1.58 | 6.96 | 6.40E-07 | 1.01E-03 |
| ion transmembrane transporter activity | molecular function | 12 | 2.54 | 4.72 | 9.93E-06 | 4.47E-03 |
| thioredoxin-disulfide reductase activity | molecular function | 2 | 0.02 | > 100 | 2.30E-04 | 4.26E-02 |
| zinc efflux active transmembrane transporter activity | molecular function | 2 | 0.02 | > 100 | 2.30E-04 | 4.03E-02 |
| zinc efflux transmembrane transporter activity | molecular function | 2 | 0.02 | 99.33 | 3.06E-04 | 4.82E-02 |
| cation efflux transmembrane transporter activity | molecular function | 2 | 0.02 | 99.33 | 3.06E-04 | 4.59E-02 |
| secondary active transmembrane transporter activity | molecular function | 8 | 1.25 | 6.39 | 4.25E-05 | 1.22E-02 |
| active transmembrane transporter activity | molecular function | 12 | 1.98 | 6.06 | 7.98E-07 | 8.39E-04 |
| active ion transmembrane transporter activity | molecular function | 10 | 1 | 9.97 | 8.85E-08 | 2.79E-04 |
| proton-transporting ATPase activity, rotational mechanism | molecular function | 3 | 0.07 | 40.64 | 7.89E-05 | 2.07E-02 |
| proton transmembrane transporter activity | molecular function | 8 | 0.81 | 9.93 | 1.89E-06 | 1.49E-03 |
| monovalent inorganic cation transmembrane transporter activity | molecular function | 8 | 1 | 8.03 | 8.66E-06 | 4.55E-03 |
| ATPase activity, coupled to transmembrane movement | molecular function | 3 | 0.07 | 40.64 | 7.89E-05 | 1.91E-02 |

| | | | | | | |
|---|---|---|---|---|---|---|
| of ions, rotational mechanism | | | | | | |
| ATP-dependent peptidase activity | molecular function | 3 | 0.11 | 27.09 | 2.39E-04 | 3.96E-02 |
| amyloplast | cellular component | 2 | 0.04 | 54.18 | 8.43E-04 | 4.19E-02 |
| plastid | cellular component | 52 | 18.43 | 2.82 | 1.71E-14 | 8.94E-12 |
| cytoplasm | cellular component | 76 | 50.08 | 1.52 | 2.05E-08 | 7.13E-06 |
| chloroplast membrane | cellular component | 9 | 0.91 | 9.86 | 4.42E-07 | 7.69E-05 |
| plastid membrane | cellular component | 9 | 0.94 | 9.61 | 5.42E-07 | 8.10E-05 |
| plastid envelope | cellular component | 14 | 2.36 | 5.93 | 1.16E-07 | 2.42E-05 |
| organelle envelope | cellular component | 16 | 3.91 | 4.09 | 1.75E-06 | 1.53E-04 |
| envelope | cellular component | 16 | 3.91 | 4.09 | 1.75E-06 | 1.67E-04 |
| chloroplast envelope | cellular component | 14 | 2.3 | 6.09 | 8.33E-08 | 2.18E-05 |
| chloroplast | cellular component | 52 | 17.05 | 3.05 | 6.30E-16 | 6.58E-13 |
| chloroplast thylakoid membrane | cellular component | 9 | 1.4 | 6.43 | 1.32E-05 | 1.06E-03 |
| chloroplast thylakoid | cellular component | 11 | 1.71 | 6.44 | 1.34E-06 | 1.56E-04 |
| plastid thylakoid | cellular component | 11 | 1.71 | 6.44 | 1.34E-06 | 1.40E-04 |
| thylakoid | cellular component | 12 | 1.94 | 6.19 | 6.46E-07 | 8.44E-05 |
| organelle subcompartment | cellular component | 11 | 3.08 | 3.57 | 2.75E-04 | 1.60E-02 |
| plastid thylakoid membrane | cellular component | 9 | 1.4 | 6.43 | 1.32E-05 | 9.82E-04 |
| thylakoid membrane | cellular component | 9 | 1.46 | 6.18 | 1.79E-05 | 1.25E-03 |
| photosynthetic membrane | cellular component | 9 | 1.46 | 6.17 | 1.82E-05 | 1.19E-03 |
| integral component of membrane | cellular component | 30 | 16.38 | 1.83 | 5.60E-04 | 3.08E-02 |
| intrinsic component of membrane | cellular component | 33 | 17.2 | 1.92 | 1.29E-04 | 7.91E-03 |
| nucleus | cellular component | 20 | 35.78 | 0.56 | 5.61E-04 | 2.93E-02 |

**Table S5. GO Term enriched in all genes with cyclic expression patterns**

| GO Term | Category | # of Genes | Expected # of Genes | Fold Enrichment | p-value | FDR |
|---|---|---|---|---|---|---|
| phenylacetate catabolic process | biological process | 3 | 0.12 | 25.2 | 1.02E-03 | 3.78E-02 |
| organic substance metabolic process | biological process | 408 | 328.39 | 1.24 | 4.49E-07 | 4.98E-05 |
| metabolic process | biological process | 495 | 381.97 | 1.3 | 5.07E-12 | 1.60E-09 |
| cellular metabolic process | biological process | 401 | 318.23 | 1.26 | 1.19E-07 | 1.55E-05 |
| cellular process | biological process | 586 | 464.75 | 1.26 | 4.71E-13 | 1.88E-10 |
| regulation of biological quality | biological process | 78 | 48.81 | 1.6 | 1.11E-04 | 5.93E-03 |
| biological regulation | biological process | 312 | 242.91 | 1.28 | 1.76E-06 | 1.59E-04 |
| response to toxic substance | biological process | 22 | 8.49 | 2.59 | 1.14E-04 | 5.97E-03 |
| response to chemical | biological process | 205 | 114.41 | 1.79 | 1.34E-15 | 8.90E-13 |
| response to stimulus | biological process | 397 | 241.52 | 1.64 | 1.70E-25 | 2.55E-22 |
| cellular response to chemical stimulus | biological process | 96 | 56.79 | 1.69 | 1.80E-06 | 1.61E-04 |
| cellular response to stimulus | biological process | 168 | 113.62 | 1.48 | 6.26E-07 | 6.36E-05 |
| carboxylic acid metabolic process | biological process | 62 | 37.03 | 1.67 | 1.88E-04 | 8.79E-03 |
| oxoacid metabolic process | biological process | 73 | 42.07 | 1.74 | 1.63E-05 | 1.13E-03 |
| organic acid metabolic process | biological process | 75 | 42.42 | 1.77 | 5.89E-06 | 4.76E-04 |
| small molecule metabolic process | biological process | 117 | 63.61 | 1.84 | 1.07E-09 | 2.01E-07 |
| photosynthesis, light harvesting in photosystem I | biological process | 15 | 0.91 | 16.43 | 3.28E-12 | 1.16E-09 |
| photosynthesis, light harvesting | biological process | 22 | 1.75 | 12.6 | 1.88E-15 | 1.12E-12 |
| photosynthesis, light reaction | biological process | 32 | 5.08 | 6.3 | 1.13E-14 | 6.17E-12 |
| generation of precursor metabolites and energy | biological process | 45 | 15.36 | 2.93 | 1.34E-09 | 2.44E-07 |
| photosynthesis | biological process | 46 | 9.25 | 4.97 | 5.09E-17 | 4.36E-14 |

| | | | | | | |
|---|---|---|---|---|---|---|
| vitamin E biosynthetic process | biological process | 5 | 0.36 | 14 | 1.20E-04 | 6.12E-03 |
| organic hydroxy compound biosynthetic process | biological process | 23 | 7.06 | 3.26 | 2.92E-06 | 2.49E-04 |
| organic hydroxy compound metabolic process | biological process | 30 | 12.1 | 2.48 | 1.76E-05 | 1.18E-03 |
| organic substance biosynthetic process | biological process | 156 | 106.71 | 1.46 | 3.22E-06 | 2.68E-04 |
| biosynthetic process | biological process | 162 | 113.22 | 1.43 | 7.16E-06 | 5.50E-04 |
| vitamin E metabolic process | biological process | 5 | 0.36 | 14 | 1.20E-04 | 6.17E-03 |
| fat-soluble vitamin metabolic process | biological process | 6 | 0.4 | 15.12 | 1.76E-05 | 1.17E-03 |
| vitamin metabolic process | biological process | 15 | 3.77 | 3.98 | 1.91E-05 | 1.25E-03 |
| fat-soluble vitamin biosynthetic process | biological process | 6 | 0.4 | 15.12 | 1.76E-05 | 1.16E-03 |
| vitamin biosynthetic process | biological process | 15 | 3.29 | 4.55 | 4.55E-06 | 3.74E-04 |
| cellular biosynthetic process | biological process | 145 | 104.41 | 1.39 | 8.97E-05 | 4.98E-03 |
| small molecule biosynthetic process | biological process | 68 | 26.75 | 2.54 | 3.47E-11 | 8.67E-09 |
| heterocycle biosynthetic process | biological process | 51 | 27.34 | 1.87 | 5.55E-05 | 3.17E-03 |
| organic cyclic compound biosynthetic process | biological process | 68 | 34.64 | 1.96 | 5.73E-07 | 6.13E-05 |
| response to low light intensity stimulus | biological process | 9 | 0.75 | 11.94 | 5.97E-07 | 6.16E-05 |
| response to light intensity | biological process | 33 | 5.79 | 5.7 | 5.12E-14 | 2.36E-11 |
| response to light stimulus | biological process | 106 | 27.86 | 3.8 | 5.01E-29 | 1.50E-25 |
| response to radiation | biological process | 106 | 28.85 | 3.67 | 6.51E-28 | 1.30E-24 |
| response to abiotic stimulus | biological process | 212 | 82.35 | 2.57 | 9.19E-35 | 5.50E-31 |
| starch catabolic process | biological process | 8 | 0.67 | 11.86 | 2.66E-06 | 2.31E-04 |
| starch metabolic process | biological process | 16 | 2.38 | 6.72 | 2.29E-08 | 3.42E-06 |
| cellular glucan metabolic process | biological process | 27 | 8.41 | 3.21 | 5.34E-07 | 5.82E-05 |
| glucan metabolic process | biological process | 27 | 8.73 | 3.09 | 1.02E-06 | 9.72E-05 |

| | | | | | | |
|---|---|---|---|---|---|---|
| polysaccharide metabolic process | biological process | 34 | 17.58 | 1.93 | 6.14E-04 | 2.44E-02 |
| cellular polysaccharide metabolic process | biological process | 29 | 11.27 | 2.57 | 1.61E-05 | 1.13E-03 |
| cellular carbohydrate metabolic process | biological process | 37 | 16.11 | 2.3 | 1.33E-05 | 9.61E-04 |
| glucan catabolic process | biological process | 11 | 2.1 | 5.23 | 2.77E-05 | 1.77E-03 |
| cellular polysaccharide catabolic process | biological process | 11 | 1.79 | 6.16 | 7.33E-06 | 5.56E-04 |
| cellular carbohydrate catabolic process | biological process | 11 | 2.82 | 3.9 | 2.79E-04 | 1.28E-02 |
| regulation of auxin biosynthetic process | biological process | 5 | 0.48 | 10.5 | 3.38E-04 | 1.50E-02 |
| regulation of biological process | biological process | 277 | 214.5 | 1.29 | 6.87E-06 | 5.41E-04 |
| regulation of cellular process | biological process | 242 | 187.95 | 1.29 | 3.82E-05 | 2.27E-03 |
| regulation of auxin metabolic process | biological process | 5 | 0.56 | 9 | 5.97E-04 | 2.40E-02 |
| xanthophyll metabolic process | biological process | 6 | 0.6 | 10.08 | 1.01E-04 | 5.50E-03 |
| cellular lipid metabolic process | biological process | 50 | 27.66 | 1.81 | 1.38E-04 | 6.77E-03 |
| lipid metabolic process | biological process | 67 | 38.61 | 1.74 | 3.24E-05 | 2.02E-03 |
| negative regulation of photomorphogenesis | biological process | 5 | 0.52 | 9.69 | 4.54E-04 | 1.91E-02 |
| regulation of photomorphogenesis | biological process | 8 | 1.27 | 6.3 | 1.14E-04 | 5.92E-03 |
| regulation of response to red or far red light | biological process | 9 | 1.67 | 5.4 | 1.19E-04 | 6.17E-03 |
| regulation of response to stimulus | biological process | 50 | 27.38 | 1.83 | 1.26E-04 | 6.34E-03 |
| regulation of post-embryonic development | biological process | 32 | 15.16 | 2.11 | 2.24E-04 | 1.03E-02 |
| negative regulation of post-embryonic development | biological process | 13 | 4.37 | 2.98 | 8.62E-04 | 3.27E-02 |
| response to high light intensity | biological process | 23 | 2.5 | 9.2 | 9.10E-14 | 3.89E-11 |
| protein-chromophore linkage | biological process | 16 | 1.75 | 9.16 | 5.61E-10 | 1.12E-07 |
| organonitrogen compound metabolic process | biological process | 253 | 190.53 | 1.33 | 2.54E-06 | 2.24E-04 |
| photosystem II repair | biological process | 5 | 0.6 | 8.4 | 7.71E-04 | 2.98E-02 |

| | | | | | | |
|---|---|---|---|---|---|---|
| protein repair | biological process | 8 | 1.15 | 6.95 | 6.32E-05 | 3.57E-03 |
| response to far red light | biological process | 16 | 2.02 | 7.91 | 3.28E-09 | 5.77E-07 |
| response to red or far red light | biological process | 43 | 8.41 | 5.11 | 2.30E-16 | 1.73E-13 |
| polyamine biosynthetic process | biological process | 5 | 0.63 | 7.87 | 9.80E-04 | 3.67E-02 |
| polyamine metabolic process | biological process | 7 | 0.95 | 7.35 | 1.36E-04 | 6.74E-03 |
| cellular biogenic amine metabolic process | biological process | 10 | 2.46 | 4.06 | 3.94E-04 | 1.71E-02 |
| cellular amine metabolic process | biological process | 10 | 2.46 | 4.06 | 3.94E-04 | 1.70E-02 |
| circadian rhythm | biological process | 34 | 4.37 | 7.79 | 7.66E-18 | 7.65E-15 |
| rhythmic process | biological process | 38 | 5.16 | 7.37 | 4.35E-19 | 5.21E-16 |
| response to red light | biological process | 20 | 2.82 | 7.1 | 1.81E-10 | 4.01E-08 |
| regulation of photosynthesis, light reaction | biological process | 7 | 1.03 | 6.78 | 2.07E-04 | 9.61E-03 |
| regulation of generation of precursor metabolites and energy | biological process | 7 | 1.27 | 5.51 | 6.11E-04 | 2.44E-02 |
| regulation of photosynthesis | biological process | 9 | 1.71 | 5.27 | 1.40E-04 | 6.80E-03 |
| response to blue light | biological process | 25 | 3.77 | 6.63 | 3.48E-12 | 1.16E-09 |
| red, far-red light phototransduction | biological process | 14 | 2.18 | 6.41 | 2.79E-07 | 3.35E-05 |
| red or far-red light signaling pathway | biological process | 16 | 3.06 | 5.24 | 4.38E-07 | 4.95E-05 |
| cellular response to red or far red light | biological process | 16 | 3.14 | 5.1 | 5.91E-07 | 6.21E-05 |
| cellular response to light stimulus | biological process | 28 | 5.24 | 5.35 | 1.54E-11 | 4.38E-09 |
| cellular response to radiation | biological process | 28 | 5.48 | 5.11 | 3.81E-11 | 9.13E-09 |
| cellular response to abiotic stimulus | biological process | 35 | 8.37 | 4.18 | 2.18E-11 | 5.68E-09 |
| cellular response to environmental stimulus | biological process | 35 | 8.37 | 4.18 | 2.18E-11 | 5.93E-09 |
| signal transduction | biological process | 98 | 67.9 | 1.44 | 4.69E-04 | 1.96E-02 |
| cell communication | biological process | 114 | 79.13 | 1.44 | 1.51E-04 | 7.25E-03 |

| | | | | | | |
|---|---|---|---|---|---|---|
| signaling | biological process | 98 | 69.25 | 1.42 | 8.45E-04 | 3.22E-02 |
| phototransduction | biological process | 14 | 2.18 | 6.41 | 2.79E-07 | 3.41E-05 |
| detection of light stimulus | biological process | 14 | 2.22 | 6.3 | 3.37E-07 | 3.88E-05 |
| detection of abiotic stimulus | biological process | 14 | 2.5 | 5.6 | 1.14E-06 | 1.07E-04 |
| detection of stimulus | biological process | 17 | 3.65 | 4.66 | 8.17E-07 | 8.16E-05 |
| detection of external stimulus | biological process | 14 | 2.5 | 5.6 | 1.14E-06 | 1.05E-04 |
| response to external stimulus | biological process | 111 | 69.37 | 1.6 | 3.14E-06 | 2.65E-04 |
| polyol biosynthetic process | biological process | 7 | 1.15 | 6.08 | 3.66E-04 | 1.60E-02 |
| alcohol biosynthetic process | biological process | 10 | 2.58 | 3.88 | 5.50E-04 | 2.22E-02 |
| alcohol metabolic process | biological process | 15 | 5.24 | 2.86 | 5.20E-04 | 2.15E-02 |
| chlorophyll biosynthetic process | biological process | 10 | 1.67 | 6 | 2.31E-05 | 1.49E-03 |
| porphyrin-containing compound biosynthetic process | biological process | 11 | 1.98 | 5.54 | 1.73E-05 | 1.18E-03 |
| cofactor biosynthetic process | biological process | 25 | 10.24 | 2.44 | 1.11E-04 | 5.91E-03 |
| cofactor metabolic process | biological process | 46 | 19.29 | 2.39 | 2.77E-07 | 3.46E-05 |
| tetrapyrrole biosynthetic process | biological process | 11 | 2.1 | 5.23 | 2.77E-05 | 1.75E-03 |
| tetrapyrrole metabolic process | biological process | 17 | 3.02 | 5.64 | 7.66E-08 | 1.02E-05 |
| aromatic compound biosynthetic process | biological process | 54 | 30.4 | 1.78 | 1.28E-04 | 6.39E-03 |
| porphyrin-containing compound metabolic process | biological process | 17 | 2.98 | 5.71 | 6.49E-08 | 8.84E-06 |
| chlorophyll metabolic process | biological process | 16 | 2.46 | 6.5 | 3.38E-08 | 4.71E-06 |
| pigment biosynthetic process | biological process | 18 | 4.92 | 3.66 | 8.27E-06 | 6.19E-04 |
| pigment metabolic process | biological process | 20 | 5.87 | 3.41 | 6.98E-06 | 5.43E-04 |
| photosystem II assembly | biological process | 6 | 1.03 | 5.82 | 1.18E-03 | 4.33E-02 |
| regulation of circadian rhythm | biological process | 12 | 2.22 | 5.4 | 9.11E-06 | 6.74E-04 |

| | | | | | | |
|---|---|---|---|---|---|---|
| monocarboxylic acid transport | biological process | 7 | 1.39 | 5.04 | 9.70E-04 | 3.65E-02 |
| organic anion transport | biological process | 18 | 7.18 | 2.51 | 6.57E-04 | 2.57E-02 |
| photomorphogenesis | biological process | 13 | 2.82 | 4.61 | 1.72E-05 | 1.19E-03 |
| response to karrikin | biological process | 23 | 5.08 | 4.53 | 1.57E-08 | 2.47E-06 |
| chloroplast organization | biological process | 39 | 9.29 | 4.2 | 1.38E-12 | 5.16E-10 |
| plastid organization | biological process | 48 | 12.02 | 3.99 | 1.95E-14 | 9.75E-12 |
| systemic acquired resistance | biological process | 9 | 2.42 | 3.72 | 1.34E-03 | 4.86E-02 |
| response to stress | biological process | 218 | 141.87 | 1.54 | 2.78E-10 | 5.74E-08 |
| aromatic amino acid family metabolic process | biological process | 11 | 3.37 | 3.26 | 1.09E-03 | 4.01E-02 |
| cell redox homeostasis | biological process | 18 | 5.67 | 3.17 | 4.61E-05 | 2.66E-03 |
| cellular homeostasis | biological process | 32 | 13.97 | 2.29 | 3.92E-05 | 2.30E-03 |
| homeostatic process | biological process | 46 | 24.84 | 1.85 | 1.42E-04 | 6.85E-03 |
| response to water deprivation | biological process | 40 | 13.77 | 2.9 | 1.37E-08 | 2.22E-06 |
| response to water | biological process | 40 | 14.09 | 2.84 | 2.42E-08 | 3.54E-06 |
| response to acid chemical | biological process | 94 | 45.16 | 2.08 | 2.31E-10 | 4.93E-08 |
| response to oxygen-containing compound | biological process | 120 | 60.88 | 1.97 | 1.25E-11 | 3.75E-09 |
| response to inorganic substance | biological process | 82 | 36.07 | 2.27 | 5.54E-11 | 1.28E-08 |
| response to reactive oxygen species | biological process | 17 | 6.07 | 2.8 | 2.89E-04 | 1.30E-02 |
| response to oxidative stress | biological process | 40 | 17.7 | 2.26 | 6.24E-06 | 4.98E-04 |
| response to wounding | biological process | 23 | 8.37 | 2.75 | 3.54E-05 | 2.14E-03 |
| response to cold | biological process | 41 | 15.99 | 2.56 | 2.33E-07 | 2.97E-05 |
| response to temperature stimulus | biological process | 60 | 23.53 | 2.55 | 5.75E-10 | 1.11E-07 |
| auxin-activated signaling pathway | biological process | 19 | 7.58 | 2.51 | 7.44E-04 | 2.89E-02 |

| | | | | | | |
|---|---|---|---|---|---|---|
| cellular response to hormone stimulus | biological process | 58 | 35.44 | 1.64 | 5.25E-04 | 2.15E-02 |
| cellular response to organic substance | biological process | 64 | 40.52 | 1.58 | 6.43E-04 | 2.53E-02 |
| response to organic substance | biological process | 130 | 74.93 | 1.74 | 3.75E-09 | 6.41E-07 |
| cellular response to endogenous stimulus | biological process | 58 | 36.19 | 1.6 | 8.21E-04 | 3.15E-02 |
| response to endogenous stimulus | biological process | 110 | 61.75 | 1.78 | 2.28E-08 | 3.50E-06 |
| response to hormone | biological process | 110 | 61.04 | 1.8 | 1.29E-08 | 2.15E-06 |
| cellular response to auxin stimulus | biological process | 21 | 8.33 | 2.52 | 3.48E-04 | 1.53E-02 |
| response to auxin | biological process | 33 | 14.92 | 2.21 | 6.61E-05 | 3.70E-03 |
| response to cadmium ion | biological process | 27 | 12.46 | 2.17 | 4.84E-04 | 2.01E-02 |
| response to metal ion | biological process | 34 | 17.06 | 1.99 | 3.23E-04 | 1.44E-02 |
| carboxylic acid biosynthetic process | biological process | 41 | 19.8 | 2.07 | 3.35E-05 | 2.05E-03 |
| organic acid biosynthetic process | biological process | 41 | 19.8 | 2.07 | 3.35E-05 | 2.07E-03 |
| response to abscisic acid | biological process | 43 | 21.23 | 2.03 | 3.74E-05 | 2.24E-03 |
| response to alcohol | biological process | 43 | 21.39 | 2.01 | 4.11E-05 | 2.39E-03 |
| response to lipid | biological process | 56 | 28.97 | 1.93 | 1.08E-05 | 7.87E-04 |
| response to salt stress | biological process | 32 | 17.1 | 1.87 | 1.30E-03 | 4.76E-02 |
| response to osmotic stress | biological process | 40 | 20.2 | 1.98 | 1.06E-04 | 5.71E-03 |
| lipid biosynthetic process | biological process | 39 | 21.15 | 1.84 | 5.46E-04 | 2.22E-02 |
| oxidation-reduction process | biological process | 107 | 59.88 | 1.79 | 3.10E-08 | 4.43E-06 |
| transmembrane transport | biological process | 57 | 33.93 | 1.68 | 2.88E-04 | 1.31E-02 |
| Unclassified | biological process | 116 | 173.94 | 0.67 | 1.01E-06 | 9.73E-05 |
| DNA metabolic process | biological process | 4 | 18.93 | 0.21 | 9.50E-05 | 5.22E-03 |
| nucleic acid metabolic process | biological process | 32 | 70.68 | 0.45 | 3.32E-07 | 3.90E-05 |

| | | | | | | |
|---|---|---|---|---|---|---|
| nucleobase-containing compound metabolic process | biological process | 49 | 86.04 | 0.57 | 1.39E-05 | 9.88E-04 |
| RNA modification | biological process | 2 | 14.37 | 0.14 | 1.52E-04 | 7.21E-03 |
| RNA metabolic process | biological process | 28 | 53.5 | 0.52 | 1.67E-04 | 7.86E-03 |
| killing of cells of other organism | biological process | 1 | 11.11 | 0.09 | 4.25E-04 | 1.81E-02 |
| cell killing | biological process | 1 | 11.11 | 0.09 | 4.25E-04 | 1.82E-02 |
| starch binding | molecular function | 5 | 0.36 | 14 | 1.20E-04 | 1.90E-02 |
| binding | molecular function | 521 | 438.4 | 1.19 | 7.42E-07 | 3.34E-04 |
| chlorophyll binding | molecular function | 16 | 1.35 | 11.86 | 2.64E-11 | 4.17E-08 |
| ion binding | molecular function | 286 | 219.62 | 1.3 | 1.91E-06 | 6.68E-04 |
| tetrapyrrole binding | molecular function | 36 | 14.84 | 2.43 | 5.41E-06 | 1.55E-03 |
| cofactor binding | molecular function | 74 | 39.65 | 1.87 | 1.25E-06 | 4.93E-04 |
| carbon-carbon lyase activity | molecular function | 18 | 4.44 | 4.05 | 2.33E-06 | 7.34E-04 |
| lyase activity | molecular function | 35 | 15.68 | 2.23 | 3.27E-05 | 7.36E-03 |
| catalytic activity | molecular function | 477 | 361.41 | 1.32 | 1.03E-12 | 3.24E-09 |
| oxidoreductase activity, acting on a sulfur group of donors | molecular function | 15 | 4.72 | 3.18 | 1.90E-04 | 2.86E-02 |
| oxidoreductase activity | molecular function | 106 | 60.44 | 1.75 | 7.72E-08 | 4.06E-05 |
| identical protein binding | molecular function | 27 | 11.43 | 2.36 | 9.01E-05 | 1.50E-02 |
| protein binding | molecular function | 271 | 196.04 | 1.38 | 3.07E-08 | 1.93E-05 |
| secondary active transmembrane transporter activity | molecular function | 34 | 14.8 | 2.3 | 2.09E-05 | 5.50E-03 |
| active transmembrane transporter activity | molecular function | 45 | 23.41 | 1.92 | 8.54E-05 | 1.50E-02 |
| transmembrane transporter activity | molecular function | 78 | 47.66 | 1.64 | 4.95E-05 | 1.04E-02 |
| transporter activity | molecular function | 81 | 50.32 | 1.61 | 5.70E-05 | 1.06E-02 |
| metal ion binding | molecular function | 172 | 126.28 | 1.36 | 5.33E-05 | 1.05E-02 |

| | | | | | | |
|---|---|---|---|---|---|---|
| cation binding | molecular function | 175 | 126.99 | 1.38 | 2.42E-05 | 5.86E-03 |
| Unclassified | molecular function | 124 | 192.31 | 0.64 | 2.17E-08 | 2.28E-05 |
| PSII associated light-harvesting complex II | cellular component | 5 | 0.28 | 18 | 5.08E-05 | 1.21E-03 |
| thylakoid light-harvesting complex | cellular component | 5 | 0.28 | 18 | 5.08E-05 | 1.18E-03 |
| chloroplast thylakoid membrane protein complex | cellular component | 8 | 0.75 | 10.61 | 5.10E-06 | 1.40E-04 |
| membrane | cellular component | 457 | 319.38 | 1.43 | 4.42E-18 | 2.56E-16 |
| cellular anatomical entity | cellular component | 1044 | 991.41 | 1.05 | 1.68E-09 | 5.49E-08 |
| chloroplast thylakoid membrane | cellular component | 80 | 16.55 | 4.83 | 5.36E-28 | 5.09E-26 |
| chloroplast thylakoid | cellular component | 99 | 20.2 | 4.9 | 6.58E-35 | 2.29E-32 |
| plastid thylakoid | cellular component | 99 | 20.2 | 4.9 | 6.58E-35 | 1.72E-32 |
| thylakoid | cellular component | 103 | 22.94 | 4.49 | 1.83E-33 | 3.83E-31 |
| intracellular | cellular component | 951 | 854.77 | 1.11 | 1.49E-13 | 6.48E-12 |
| plastid | cellular component | 437 | 217.99 | 2 | 1.10E-49 | 1.15E-46 |
| intracellular membrane-bounded organelle | cellular component | 877 | 774.89 | 1.13 | 3.67E-12 | 1.47E-10 |
| intracellular organelle | cellular component | 886 | 787.35 | 1.13 | 1.10E-11 | 3.95E-10 |
| organelle | cellular component | 889 | 791.32 | 1.12 | 1.17E-11 | 4.09E-10 |
| membrane-bounded organelle | cellular component | 887 | 783.06 | 1.13 | 8.78E-13 | 3.67E-11 |
| cytoplasm | cellular component | 776 | 592.22 | 1.31 | 4.46E-29 | 5.83E-27 |
| organelle subcompartment | cellular component | 110 | 36.47 | 3.02 | 1.19E-22 | 8.29E-21 |
| chloroplast | cellular component | 408 | 201.68 | 2.02 | 1.70E-46 | 8.89E-44 |
| plastid thylakoid membrane | cellular component | 80 | 16.55 | 4.83 | 5.36E-28 | 4.67E-26 |
| thylakoid membrane | cellular component | 82 | 17.22 | 4.76 | 2.79E-28 | 3.24E-26 |
| photosynthetic membrane | cellular component | 82 | 17.26 | 4.75 | 3.20E-28 | 3.34E-26 |

| | | | | | | |
|---|---|---|---|---|---|---|
| light-harvesting complex | cellular component | 5 | 0.28 | 18 | 5.08E-05 | 1.15E-03 |
| amyloplast | cellular component | 5 | 0.44 | 11.45 | 2.47E-04 | 5.15E-03 |
| plastoglobule | cellular component | 30 | 3.14 | 9.57 | 6.48E-18 | 3.56E-16 |
| chloroplast stroma | cellular component | 109 | 30.68 | 3.55 | 1.32E-27 | 9.88E-26 |
| plastid stroma | cellular component | 110 | 31.11 | 3.54 | 1.08E-27 | 8.72E-26 |
| photosystem I reaction center | cellular component | 4 | 0.44 | 9.16 | 2.06E-03 | 3.58E-02 |
| photosystem I | cellular component | 18 | 1.63 | 11.06 | 3.79E-12 | 1.47E-10 |
| photosystem | cellular component | 27 | 3.61 | 7.48 | 4.25E-14 | 1.93E-12 |
| photosystem II | cellular component | 19 | 2.62 | 7.25 | 3.78E-10 | 1.27E-08 |
| chloroplast inner membrane | cellular component | 16 | 3.49 | 4.58 | 2.06E-06 | 6.15E-05 |
| plastid inner membrane | cellular component | 16 | 3.61 | 4.43 | 3.02E-06 | 8.54E-05 |
| plastid membrane | cellular component | 47 | 11.07 | 4.24 | 4.84E-15 | 2.30E-13 |
| organelle membrane | cellular component | 135 | 87.11 | 1.55 | 1.07E-06 | 3.39E-05 |
| plastid envelope | cellular component | 109 | 27.94 | 3.9 | 1.16E-30 | 2.01E-28 |
| organelle envelope | cellular component | 121 | 46.23 | 2.62 | 3.76E-20 | 2.31E-18 |
| envelope | cellular component | 121 | 46.23 | 2.62 | 3.76E-20 | 2.46E-18 |
| chloroplast membrane | cellular component | 47 | 10.79 | 4.35 | 2.08E-15 | 1.09E-13 |
| chloroplast envelope | cellular component | 107 | 27.18 | 3.94 | 2.14E-30 | 3.20E-28 |
| chloroplast thylakoid lumen | cellular component | 9 | 2.3 | 3.91 | 9.74E-04 | 1.82E-02 |
| plastid thylakoid lumen | cellular component | 9 | 2.3 | 3.91 | 9.74E-04 | 1.79E-02 |
| thylakoid lumen | cellular component | 12 | 2.9 | 4.14 | 9.08E-05 | 2.02E-03 |
| secretory vesicle | cellular component | 16 | 6.59 | 2.43 | 2.54E-03 | 4.28E-02 |
| peroxisome | cellular component | 28 | 12.78 | 2.19 | 3.32E-04 | 6.68E-03 |

| | | | | | | |
|---|---|---|---|---|---|---|
| microbody | cellular component | 28 | 12.78 | 2.19 | 3.32E-04 | 6.81E-03 |
| vacuolar membrane | cellular component | 44 | 24.92 | 1.77 | 6.83E-04 | 1.32E-02 |
| whole membrane | cellular component | 61 | 38.26 | 1.59 | 6.27E-04 | 1.24E-02 |
| vacuole | cellular component | 98 | 45.24 | 2.17 | 1.00E-11 | 3.73E-10 |
| bounding membrane of organelle | cellular component | 75 | 51.79 | 1.45 | 2.39E-03 | 4.09E-02 |
| cytosol | cellular component | 224 | 130.01 | 1.72 | 2.92E-15 | 1.45E-13 |
| plasma membrane | cellular component | 204 | 150.56 | 1.35 | 1.01E-05 | 2.45E-04 |
| cell periphery | cellular component | 231 | 173.15 | 1.33 | 6.51E-06 | 1.74E-04 |
| integral component of membrane | cellular component | 257 | 193.74 | 1.33 | 2.40E-06 | 6.97E-05 |
| intrinsic component of membrane | cellular component | 269 | 203.46 | 1.32 | 1.41E-06 | 4.34E-05 |
| ribosomal subunit | cellular component | 2 | 12.34 | 0.16 | 8.91E-04 | 1.69E-02 |
| ribonucleoprotein complex | cellular component | 10 | 28.49 | 0.35 | 1.18E-04 | 2.52E-03 |
| mitochondrial protein complex | cellular component | 1 | 9.37 | 0.11 | 1.81E-03 | 3.21E-02 |
| nuclear chromosome | cellular component | 0 | 7.78 | < 0.01 | 1.02E-03 | 1.84E-02 |
| nuclear lumen | cellular component | 20 | 43.26 | 0.46 | 1.16E-04 | 2.52E-03 |
| intracellular organelle lumen | cellular component | 23 | 52.46 | 0.44 | 7.49E-06 | 1.96E-04 |
| organelle lumen | cellular component | 23 | 52.46 | 0.44 | 7.49E-06 | 1.86E-04 |
| membrane-enclosed lumen | cellular component | 23 | 52.46 | 0.44 | 7.49E-06 | 1.91E-04 |

**Table S6. Parameter values for yearly daylength models**

| Location | a | b | c |
|---|---|---|---|
| Oslo | 0.11 | 0.76 | 0.437 |
| Paria | 0.1 | 0.12 | 0.437 |
| Boston | 0.11 | 0.4 | 0.434 |

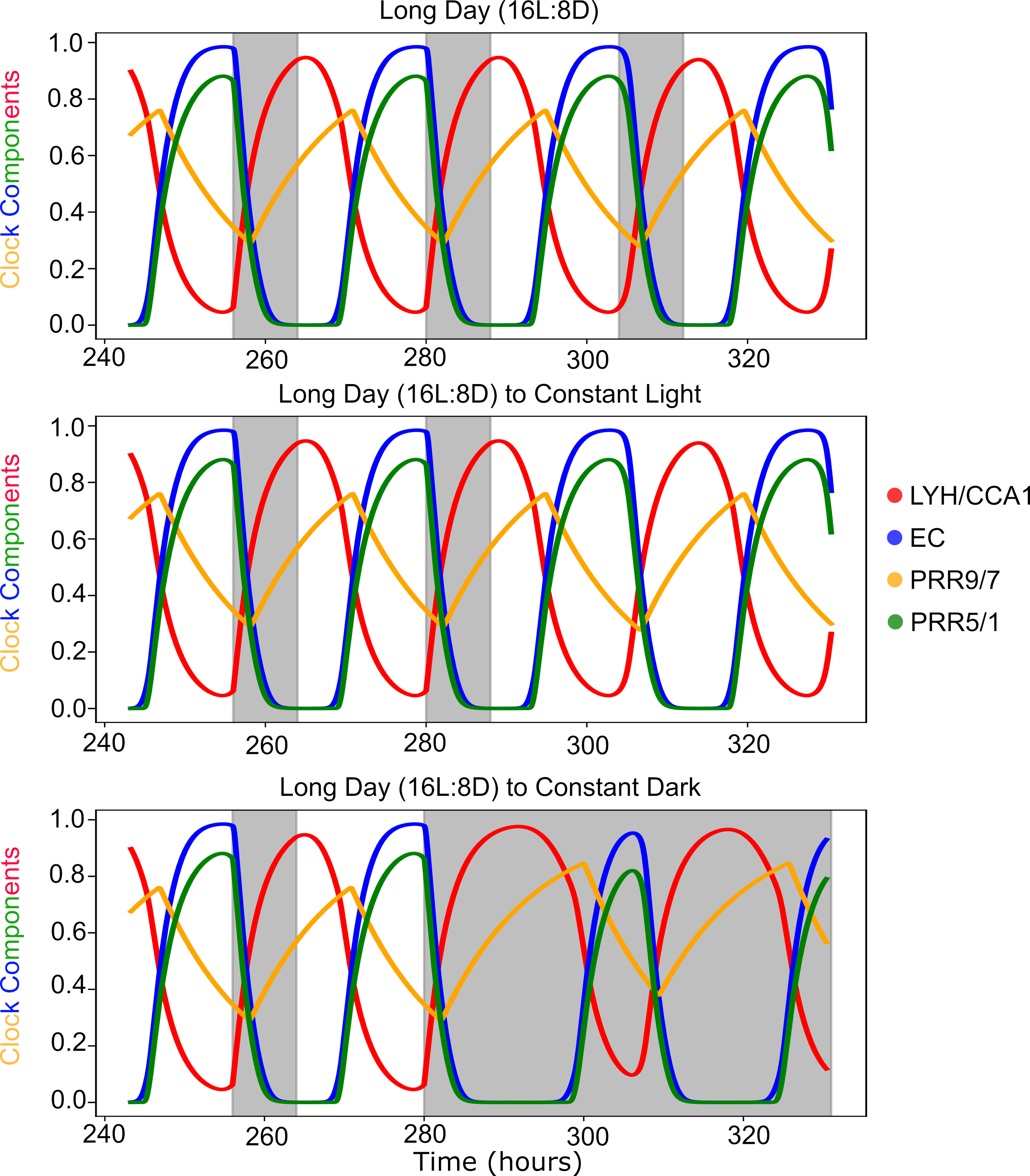

Long Day (16L:8D)
Long Day (16L:8D) to Constant Light
Long Day (16L:8D) to Constant Dark
Clock Components
Time (hours)
LYH/CCA1
EC
PRR9/7
PRR5/1

**Fig. S1. The cyclic behavior of the clock model**. The behavior of the clock modules under constant long days (16L:8D), transition from long-days to constant light and transition from long-days to constant dark. The four components of the clock are indicated by color (red = LHY/CCA1 = C1, orange = PRR9/7 = C3, green = PRR5/1 = C4, blue = EC = C2). The grey shaded area indicates the night phase of the light-dark cycle.

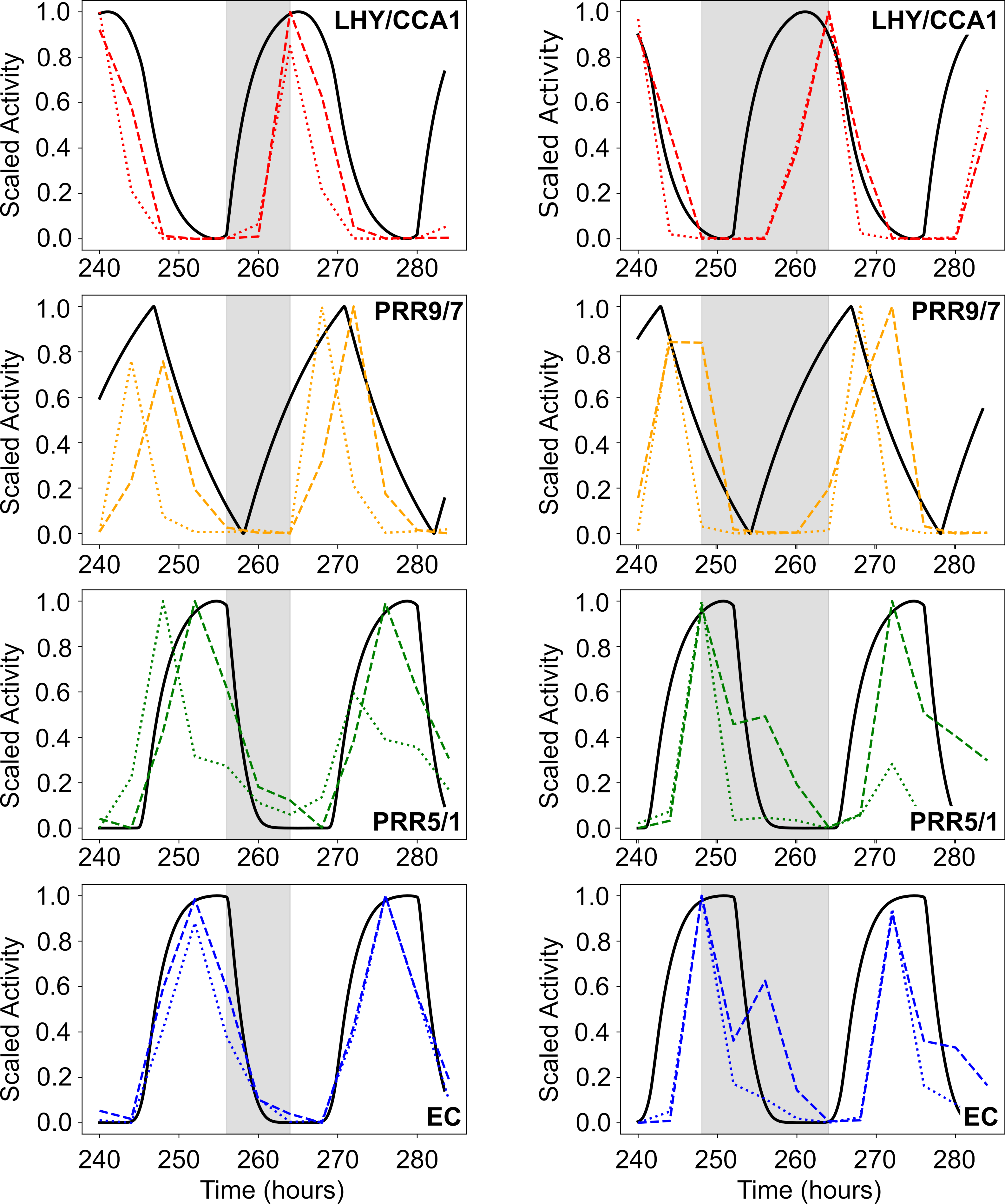

LHY/CCA1
PRR9/7
PRR5/1
EC
Scaled Activity
Time (hours)

**Fig. S2. Comparison of the clock model to the expression of circadian factors from the Diurnal Database**. Simulated activity (black) is plotted against normalized expression of circadian factors that are indicated by color (red = LHY and CCA1, orange = PRR9 and PPR7, green = PRR5 and PRR1, blue = ELF4 and LUX) (ref 28). Long day (16L:8D) data/simulations are on the right, and short day (8L:16D) data/simulations are on the left. The grey shaded area indicates the night phase of the light-dark cycle. Experimental and model values were scaled to [0, 1] based on the minima and maxima.

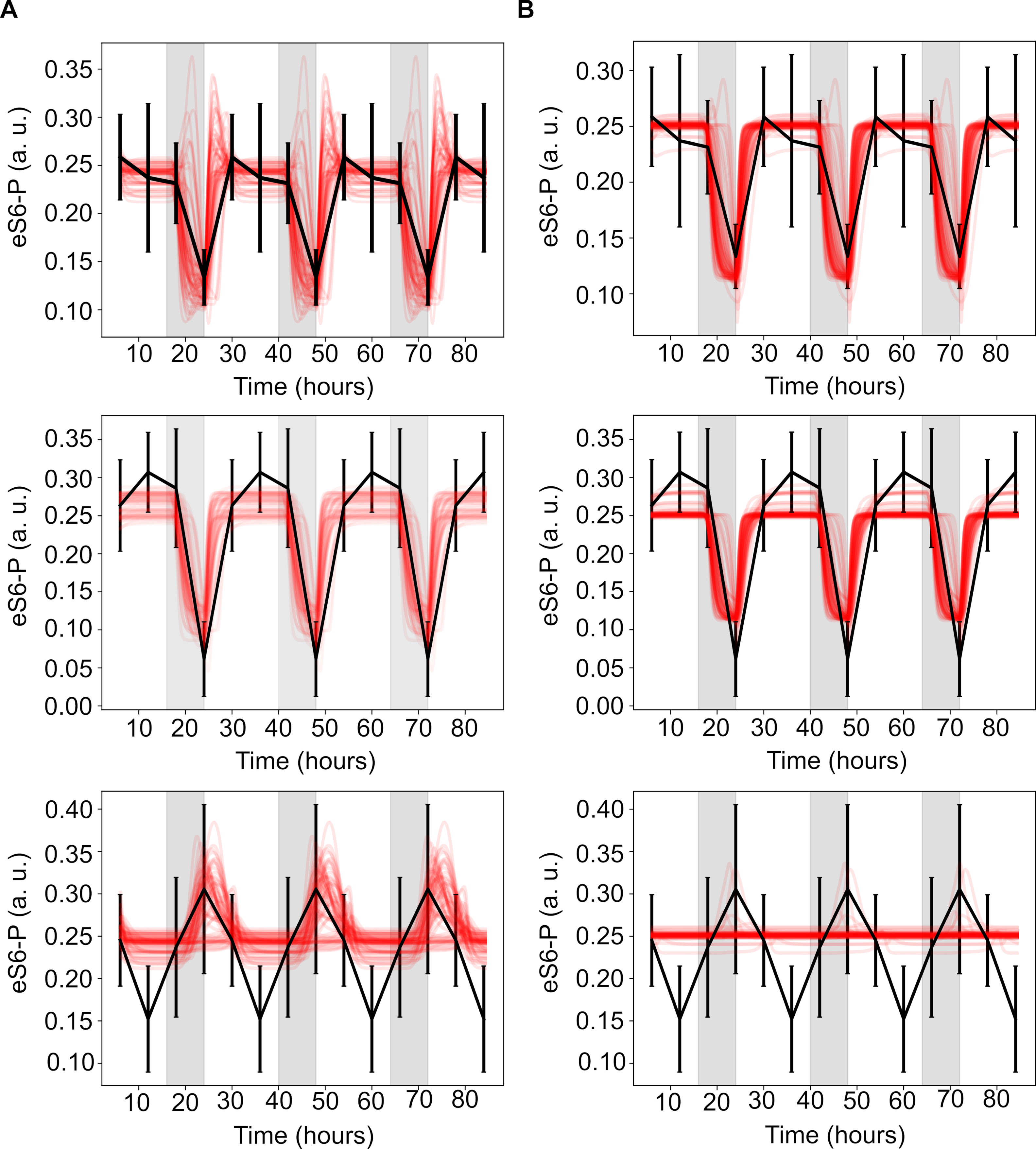
A
B
eS6-P (a. u.)
Time (hours)

**Fig. S3. Performance of the eS6-P model using alternatively optimized parameters sets. (A)** Behavior of the eS6-P model using the top 70 (5%) parameter sets among the 1400 optimization runs under long-day (top), CCA1-overexpression (middle) and constant light (bottom) conditions. The black curve indicates the experimental data to which the models were fit, and each red curve is a trajectory generated from a parameter set. The grey shaded area indicates the night phase of the light-dark cycle or subjective night in the case of constant light. Note that these models are consistent with the model in Fig. 1E in terms of overall score and specific features, including an early day peak under a regular long-day and cyclic behavior under constant light. **(B)** Behavior of the eS6-P model using the next 70 optimized parameter sets under long-day (top), CCA1-overexpression (middle) and constant light (bottom) conditions. The black curve indicates the experimental data to which the models were fit, and each red curve is a trajectory generated from a parameter set. The grey shaded area indicates the night phase of the light-dark cycle or subjective night in the case of constant light.

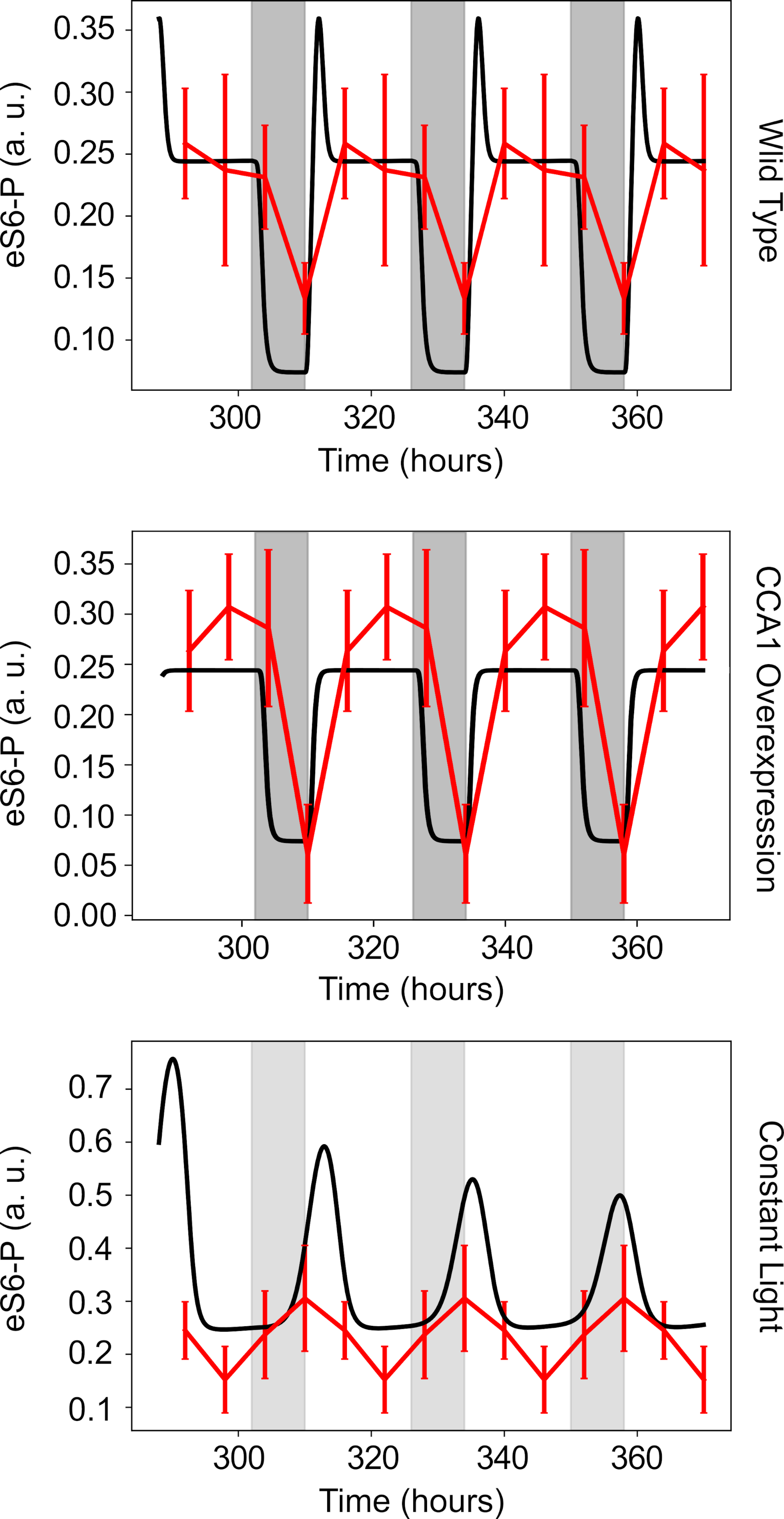
eS6-P (a. u.)
0.35
0.30
0.25
0.20
0.15
0.10
300
320
340
360
Time (hours)
Wild Type
eS6-P (a. u.)
0.35
0.30
0.25
0.20
0.15
0.10
0.05
0.00
300
320
340
360
Time (hours)
CCA1 Overexpression
eS6-P (a. u.)
0.7
0.6
0.5
0.4
0.3
0.2
0.1
300
320
340
360
Time (hours)
Constant Light

**Fig. S4. Simulations based on a model of eS6-P using a detailed circadian clock model from De Caluwe et al.** (ref 25). Trajectories from simulations (black) are compared to observations from long-day (top), CCA1-overexpression (middle) and constant light (bottom) conditions. Circadian time is measured in hours relative to subjective dawn and shaded grey regions indicate periods of subjective night. See Methods for details of this clock model.

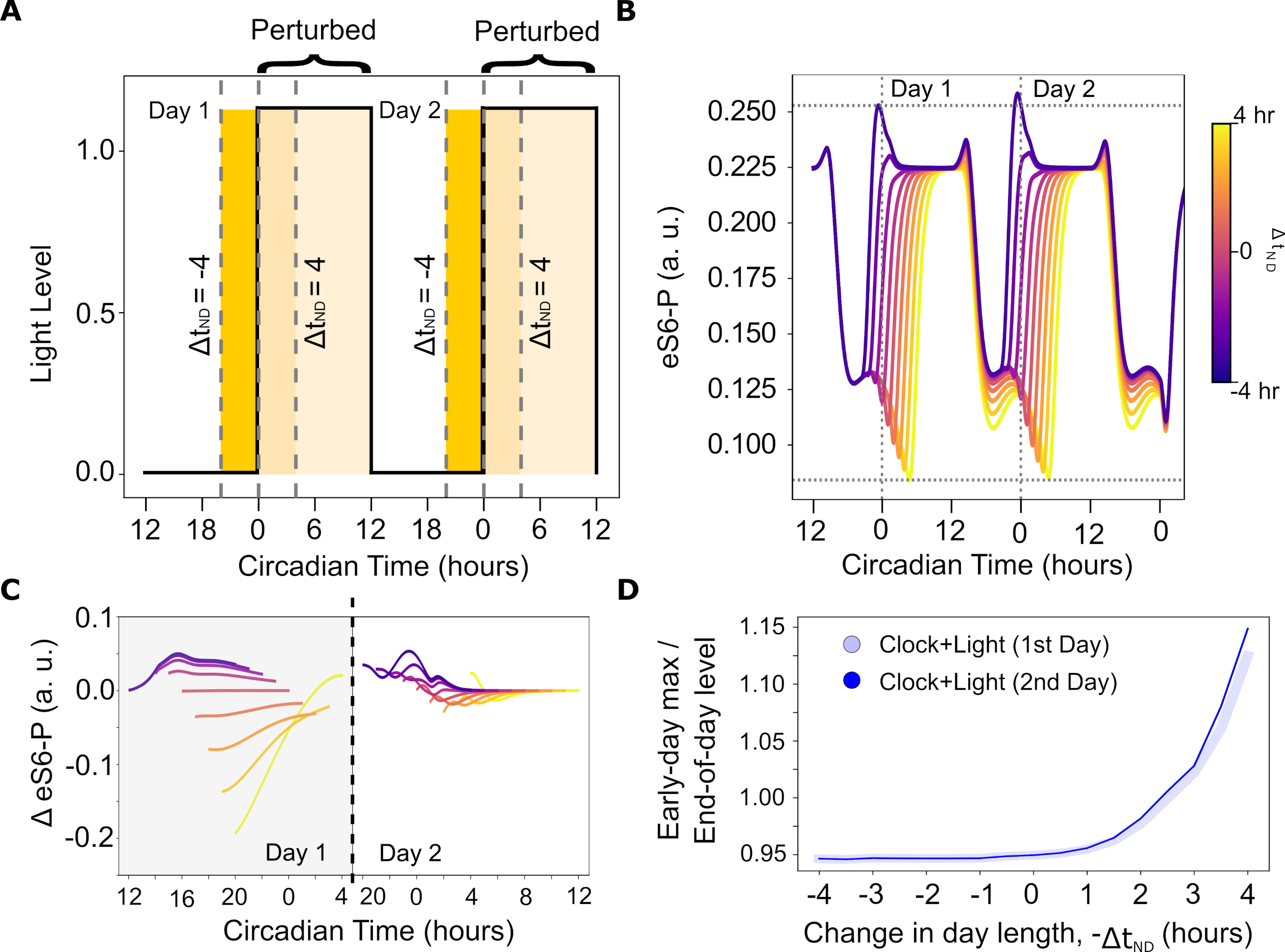

A
Perturbed
Perturbed
Day 1
Day 2
Light Level
$\Delta t_{ND} = -4$
$\Delta t_{ND} = 4$
$\Delta t_{ND} = -4$
$\Delta t_{ND} = 4$
1.0
0.5
0.0
12 18 0 6 12 18 0 6 12
Circadian Time (hours)
B
Day 1
Day 2
eS6-P (a. u.)
0.250
0.225
0.200
0.175
0.150
0.125
0.100
12 0 12 0 12 0
Circadian Time (hours)
4 hr
0
-4 hr
$\Delta t_{ND}$
C
0.1
0.0
-0.1
-0.2
Δ eS6-P (a. u.)
Day 1
Day 2
12 16 20 0 4 20 0 4 8 12
Circadian Time (hours)
D
Early-day max /
End-of-day level
Clock+Light (1st Day)
Clock+Light (2nd Day)
1.15
1.10
1.05
1.00
0.95
-4 -3 -2 -1 0 1 2 3 4
Change in day length, $-\Delta t_{ND}$ (hours)

**Fig. S5. Response of eS6-P to variation in the night to day transition time over consecutive days. (A)** Diagram of perturbations of the night to day transition and the effect on length of the day relative to a normal 12L:12D day for two consecutive perturbed days. The extent of the day is shown by yellow shaded regions and the extent of the change in day length is shown by differential shading. **(B)** Early day behavior of eS6-P on the second day in response to varying the night to day transition time from $\Delta t_{ND}$ = -4 (purple) to $\Delta t_{ND}$ = 4 (yellow) in 1-hour increments for two consecutive days. Each model was measured for 8 hours after dawn as the shortest day is 8 hours. **(C)** The difference in eS6-P predicted by the model around the first and second dawn. The difference over the last eight hours before dusk (grey) is shown on the left and the difference in the first eight hours after dawn (white) is shown on the right. **(D)** Early day peak metric (ratio of the early day maximum of eS6-P to the eS6-P levels at the dusk) of eS6-P across different degrees of dawn variation on two consecutive days. Thin solid lines show early eS6-P response on the second day, while thicker transparent lines show early eS6-P response on the first day.

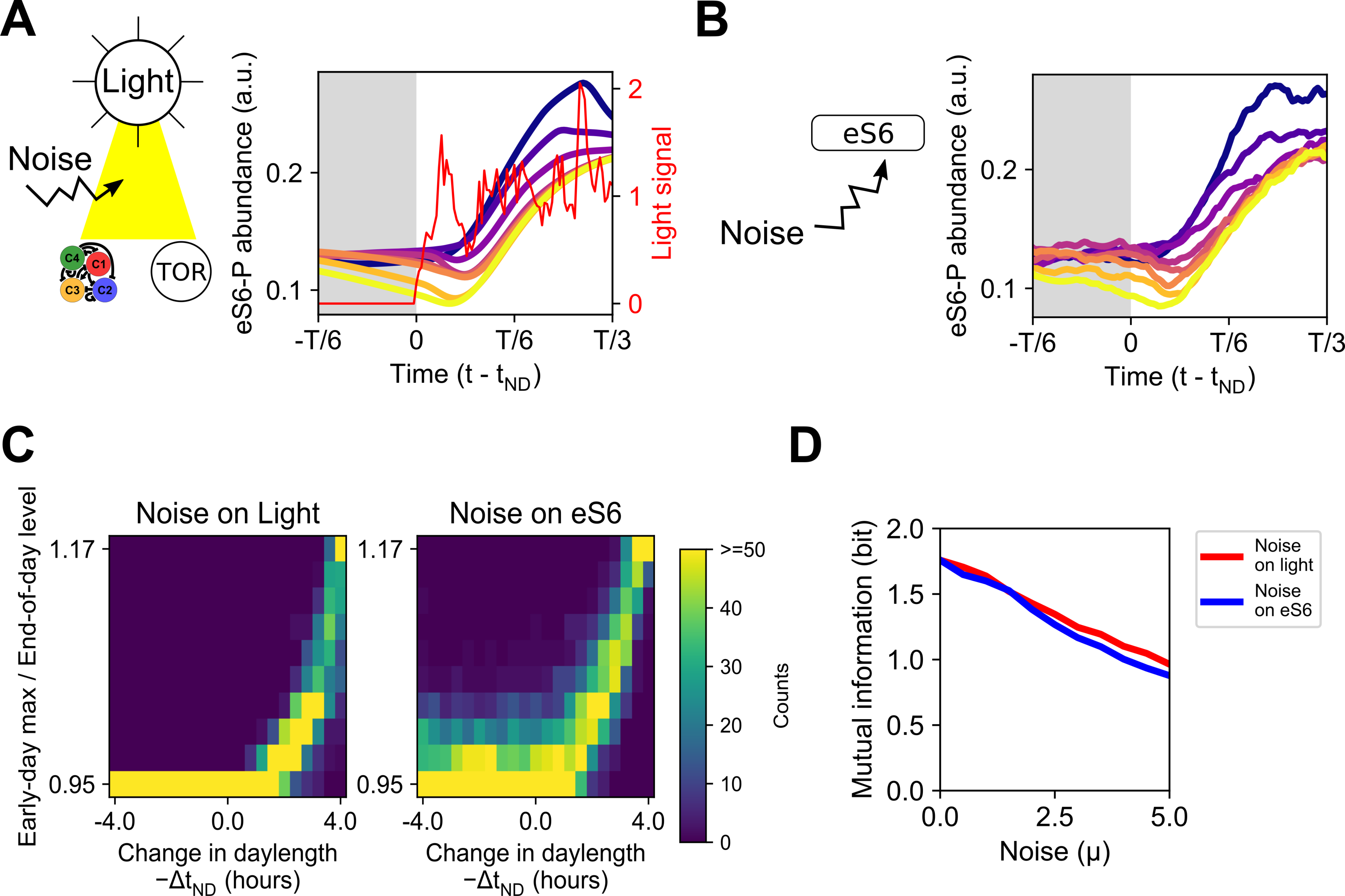

A
Light
Noise
C4
C1
C3
C2
TOR
eS6-P abundance (a.u.)
Light signal
0.2
0.1
2
1
0
-T/6
0
T/6
T/3
Time (t - t_ND)
B
eS6
Noise
eS6-P abundance (a.u.)
0.2
0.1
-T/6
0
T/6
T/3
Time (t - t_ND)
C
Noise on Light
Noise on eS6
Early-day max / End-of-day level
1.17
0.95
-4.0
0.0
4.0
Change in daylength
−Δt_ND (hours)
>=50
40
30
20
10
0
Counts
D
Mutual information (bit)
2.0
1.5
1.0
0.5
0.0
2.5
5.0
Noise (μ)
Noise on light
Noise on eS6

**Fig. S6. Detection of daylength variations with early-day eS6-P response in the presence of light fluctuations and concentration fluctuations**. **(A)** Five representative trajectories from simulations of the Clock+Light model (Fig. 1E). For each simulation, the system first reached the steady state under 12-hour light and 12-hour dark condition. Next, a perturbation of the time at which the light is turned on was performed at the dawn ($\Delta t_{ND}$, positive perturbation represents postponed dawn time). Trajectories were aligned at the actual dawn. To model fluctuations of light, a white noise term with an amplitude parameter $\mu$=3 was added to the differential equation describing the light signal (see Methods). Red curve shows a representative light signal is shown. **(B)** Five representative trajectories from simulations with noise term on eS6-P ($\mu$=3). **(C)** Contingency tables summarizing the relationship between daylength variations and the early eS6-P levels under the conditions in **(A)** and **(B)**. 200 stochastic simulations were performed for each daylength. eS6-P responses were categorized into 10 bins, and 21 daylength variations were tested. **(D)** Mutual information between daylength variations (represented by $\Delta t_{ND}$) and the early eS6-P levels. Multiple noise amplitudes were analyzed for the two conditions indicated.
